# Congenital heart defects in Down syndrome are caused by increased dosage of DYRK1A

**DOI:** 10.1101/2023.09.18.558244

**Authors:** Eva Lana-Elola, Rifdat Aoidi, Miriam Llorian, Dorota Gibbins, Callan Buechsenschuetz, Claudio Bussi, Helen Flynn, Tegan Gilmore, Sheona Watson-Scales, Marie Haugsten Hansen, Darryl Hayward, Ok-Ryul Song, Véronique Brault, Yann Herault, Emmanuel Deau, Laurent Meijer, Ambrosius P. Snijders, Maximiliano Gutierrez, Elizabeth M. C. Fisher, Victor L. J. Tybulewicz

## Abstract

Down syndrome (DS), trisomy 21, is a gene dosage disorder which results in multiple phenotypes including congenital heart defects (CHD). This clinically important pathology is caused by a third copy of one or more of the ∼230 genes on human chromosome 21 (Hsa21), but the identity of the causative dosage-sensitive genes is unknown and hence pathological mechanisms remain obscure. We show that embryonic hearts from human fetuses with DS and mouse models of DS have reduced expression of mitochondrial respiration and cell proliferation genes correlating with CHD. Using systematic genetic mapping, we determine that three copies of the *Dyrk1a* gene, encoding a serine/threonine protein kinase, are required to cause CHD. Reducing *Dyrk1a* copy number from three to two reverses defects in proliferation and mitochondrial respiration in embryonic cardiomyocytes and rescues septation defects in DS hearts. Furthermore, treatment of pregnant mice with a DYRK1A inhibitor developed for clinical use partially reduces the incidence of CHD among Dp1Tyb embryos. Thus, increased dosage of DYRK1A is required to impair mitochondrial function and cause CHD in DS, revealing a therapeutic target for this common human condition.

**One Sentence Summary:** Increased dosage of DYRK1A causes mitochondrial dysfunction and congenital heart defects in Down syndrome and is ameliorated in utero by a drug.

## Introduction

Down syndrome (DS), trisomy of human chromosome 21 (Hsa21), is a common human condition resulting in many different phenotypes, including learning and memory deficits, craniofacial alterations, early-onset Alzheimer’s disease and congenital heart defects (CHD) (*1*). DS is a gene dosage disorder with an extra copy of one or more of the genes on Hsa21 resulting in the different phenotypes. However, the identities of these causative dosage-sensitive genes remain largely unknown (*1, 2*). Discovery of such causative genes would facilitate study of the pathological mechanisms, paving the way for therapies, which are lacking for most DS clinical conditions.

With an estimated prevalence of around 1 in 800 births, DS is the most common genetic cause of CHD, with around 50% of babies with DS presenting with cardiac defects at birth (*3, 4*). Typically, these are ventricular and atrioventricular septal defects (VSDs and AVSDs) and malformations of the outflow tract (*3–5*). The most severe defects, such as AVSDs, often require surgical intervention in the first years after birth, resulting in significant morbidity and mortality (*6*). Several proteins encoded by Hsa21 genes have been proposed as candidates for the CHD. These include DSCAM and JAM2 adhesion molecules, COL6A1, COL6A2 and COL18A1 collagens, ADAMTS1 and ADAMTS5 metallopeptidases, the DYRK1A kinase, RCAN1, an inhibitor of calcineurin phosphatase, and SYNJ1, a regulator of endosomes (*7*). However, there has been no direct genetic demonstration of causality for any of these genes in studies that modulate gene dosage from 3 to 2 (*2*). Hence pathological mechanisms underlying this clinically important condition are not understood.

Hsa21 is orthologous to three regions of the mouse genome on mouse chromosome 10 (Mmu10), Mmu16 and Mmu17, with the largest of these being on Mmu16 (*2*). In previous work we generated the Dp1Tyb mouse model of DS that has an extra copy of the entire 23 Mb Hsa21-orthologous region of Mmu16 containing 145 coding genes in a tandem duplication, thereby genetically recapitulating trisomy of around 62% of Hsa21 genes (*8*). Dp1Tyb mice have a broad range of DS-like phenotypes (*8, 9*). Importantly, these mice have CHD similar to those seen in humans with DS, with ∼50% of embryonic day 14.5 (E14.5) Dp1Tyb embryos showing VSDs, AVSDs and outflow tract defects (*8*). Furthermore, making use of a series of mouse strains with an extra copy of shorter segments of the Hsa21-orthologous region of Mmu16, we were able to show that an extra copy of just 39 coding genes in Dp3Tyb mice is sufficient to cause CHD, and that this region must contain at least two causative genes (*8*).

Here we use transcriptomics of human DS fetal hearts and mouse embryonic hearts from models of DS to discover that reduced expression of mitochondrial respiration and cell proliferation genes correlates with CHD. Using systematic genetic mapping we demonstrate that one of the causative genes for CHD in DS is *Dyrk1a*, which encodes the DYRK1A serine/threonine kinase. We show that increased dosage of *Dyrk1a* results in impaired cell proliferation and mitochondrial respiration of cardiomyocytes and is required to cause CHD in DS. Finally, we demonstrate that treatment of pregnant mice with a DYRK1A inhibitor partially reduces the incidence of CHD among Dp1Tyb embryos.

## Results

### Transcriptomic similarities in embryonic hearts from human fetuses with DS and mouse models of DS

To discover biochemical pathways that are altered in DS and contribute to the pathology of the cardiac defects, we used RNA sequencing (RNAseq) to analyze the transcriptome of DS fetal hearts and age- and sex-matched euploid controls (Table S1). Differential gene expression analysis showed that DS hearts had, as expected, upregulated expression of many Hsa21 genes (Figure 1A, Table S2). In addition, expression of a further 99 genes was significantly altered and hierarchical clustering demonstrated that the transcriptomes of DS and euploid fetal hearts are distinct (Figures 1A, B).

**Figure 1.**
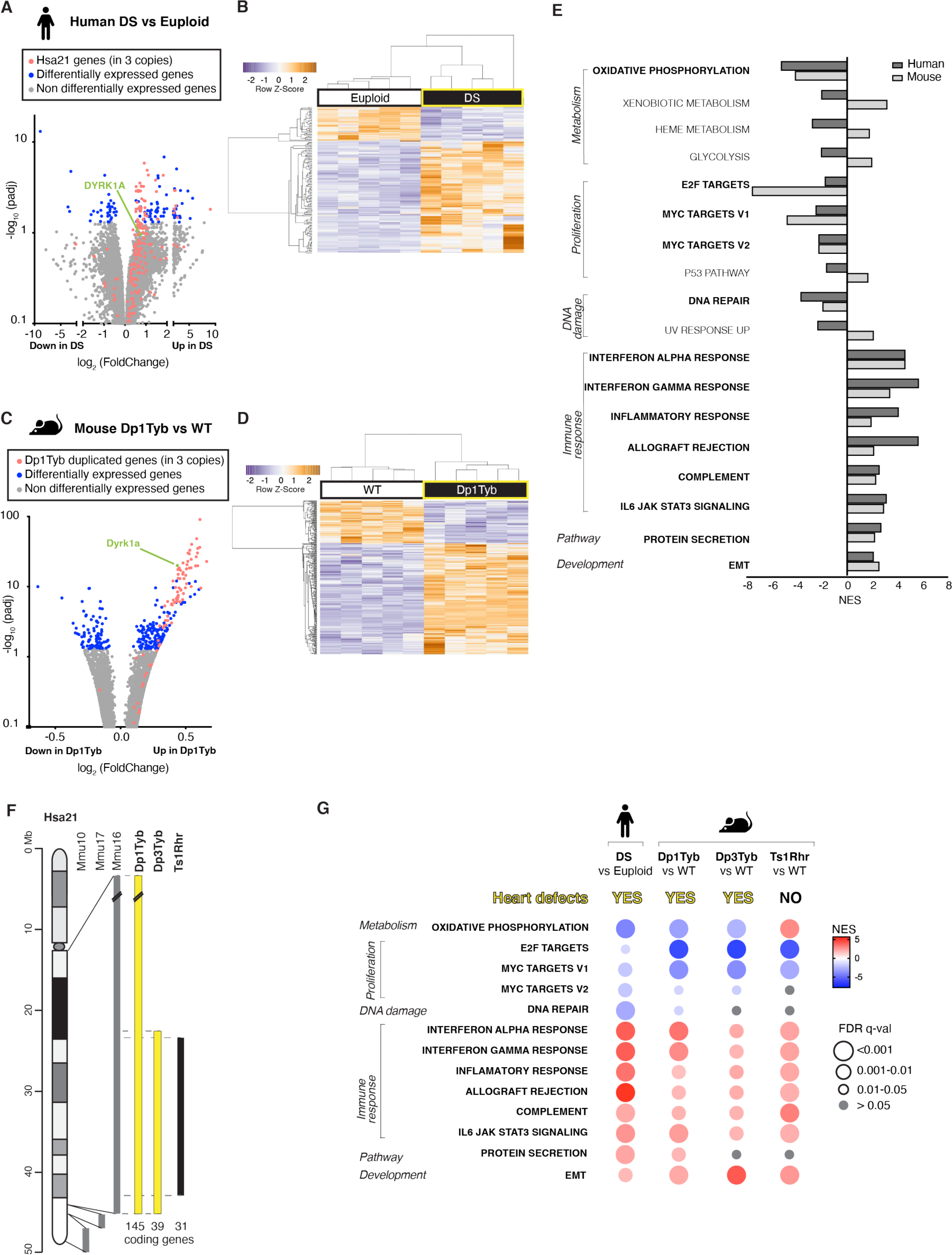
Transcriptomic similarities in embryonic hearts from human fetuses with DS and mouse models of DS. (**A, B**) RNAseq analysis of human DS and euploid embryonic hearts (13-14 post-conception weeks, n=5), showing (A) a volcano plot of fold-change in gene expression (DS v euploid) against adjusted P-value for significance of the difference, indicating Hsa21 genes and differentially expressed genes, (B) unsupervised hierarchical clustering of the 10 samples showing heatmap of differentially expressed genes. *DYRK1A* is indicated on the volcano plot. (**C, D)** RNAseq analysis of E13.5 hearts from WT and Dp1Tyb embryos (n=5) showing (C) a volcano plot as in A indicating genes in 3 copies in Dp1Tyb mice and (D) differentially expressed genes, and hierarchical clustering of the samples as in B. *Dyrk1a* is indicated on the volcano plot. (**E**) Hallmark gene sets from the Molecular Signatures Database that are significantly enriched or depleted (GSEA, 5% FDR) in both human DS and Dp1Tyb mouse hearts; NES, normalized enrichment scores. Gene sets showing the same direction of change in human and mouse data are indicated in bold. (**F**) Map of Hsa21 (length in Mb) showing cytogenetic bands and regions of orthology to Mmu10, Mmu17 and Mmu16 (grey) and indicating regions of Mmu16 that are duplicated in mouse strains (bold) that show CHD (yellow) and that do not (black); numbers of coding genes indicated below duplicated regions. (**G**) Comparison of dysregulated gene sets determined by GSEA of RNAseq data from human DS vs euploid embryonic hearts and in hearts from Dp1Tyb, Dp3Tyb and Ts1Rhr mouse embryos compared to WT controls. All show heart defects except Ts1Rhr mice. Colors and sizes of circles indicate NES and FDR q-value, respectively. Sample numbers: n=5 embryonic hearts (A, C).

To investigate further, we made use of the Dp1Tyb mouse model of DS, which has CHD similar to those seen in humans with DS (*8*). Focusing on E13.5, the day before septation is completed, RNAseq analysis of Dp1Tyb and wild-type (WT) control hearts showed that the genes present in three copies in Dp1Tyb mice are increased in expression by about 1.5-fold, as expected (Figure 1C, Figure S1A, Table S3). In addition, a further 242 genes were significantly altered in expression in Dp1Tyb embryonic hearts, and hierarchical clustering showed that Dp1Tyb and WT embryonic hearts are transcriptionally distinct (Figure 1D). Gene set expression analysis (GSEA) (*10*) revealed significant changes in multiple pathways in human DS and mouse Dp1Tyb embryonic hearts (Figure 1E). Most of these were significantly altered in the same direction in both species, including decreased oxidative phosphorylation and proliferation, and increased immune responses and epithelial to mesenchymal transition (EMT), showing that the transcriptional changes in Dp1Tyb embryonic hearts resemble those in human DS hearts (Figure 1E).

### Decreased expression of oxidative phosphorylation and cell proliferation genes correlates with CHD

To identify which pathways were most likely to be involved in CHD pathogenesis, we extended the RNAseq analysis to two further mouse strains: Dp3Tyb and Ts1Rhr. Dp3Tyb embryos show CHD similar to those in Dp1Tyb mice and DS, but this strain has an extra copy of just 39 protein-coding genes, entirely contained within the large duplication in Dp1Tyb mice, thus demonstrating that these 39 genes are sufficient to cause CHD (Figure 1F) (*8*). In contrast, Ts1Rhr embryos have an extra copy of a slightly shorter region containing just 31 genes and do not show CHD (*8*). Comparison of the transcriptional changes in Dp3Tyb and Ts1Rhr embryonic hearts with those of Dp1Tyb and DS hearts, showed that the changes in DNA repair and protein secretion pathways were not seen in either Dp3Tyb or Ts1Rhr hearts and these were not considered further. Increased expression of immune response and EMT pathways was seen in both Dp3Tyb and Ts1Rhr hearts, demonstrating that these changes are not sufficient to cause CHD (Figure 1G, Tables S4 and S5). In contrast, decreased expression of oxidative phosphorylation genes correlated with CHD, being seen in human DS fetal hearts and in Dp1Tyb and Dp3Tyb embryonic hearts, but not in Ts1Rhr hearts (Figure 1G, Figure S1B, C). Decreased expression of the 3 proliferation gene sets was seen in Dp3Tyb hearts, but only 2 of these were also decreased in Ts1Rhr hearts. These results suggest that impaired oxidative phosphorylation, and potentially proliferation are important contributors to the etiology of CHD.

### scRNAseq reveals similar gene expression changes across different cell types in Dp1Tyb embryonic hearts

The developing mammalian heart contains many different cell types. The RNAseq analysis identified dysregulated pathways but was not able to specify which cell type is perturbed in Dp1Tyb hearts. To address this, we performed single cell RNAseq (scRNAseq) of WT and Dp1Tyb E13.5 hearts and identified 14 clusters which were assigned to individual cell types based on expression of marker genes (Figures 2A, B) (*11*). Many of the expected cell types were detected, including several sub-types of cardiomyocytes, endocardial cells, epicardial cells, fibroblasts, vascular endothelial cells, smooth muscle cells and macrophages (*11*). Comparison of the frequencies of each cell type showed that while all cell types were detected in each genotype (Figure S2), Dp1Tyb hearts had reduced frequencies of ventricular and atrial cardiomyocytes and increased frequencies of fibroblasts and vascular endothelial cells compared to WT (Figure 2C).

**Figure 2.**
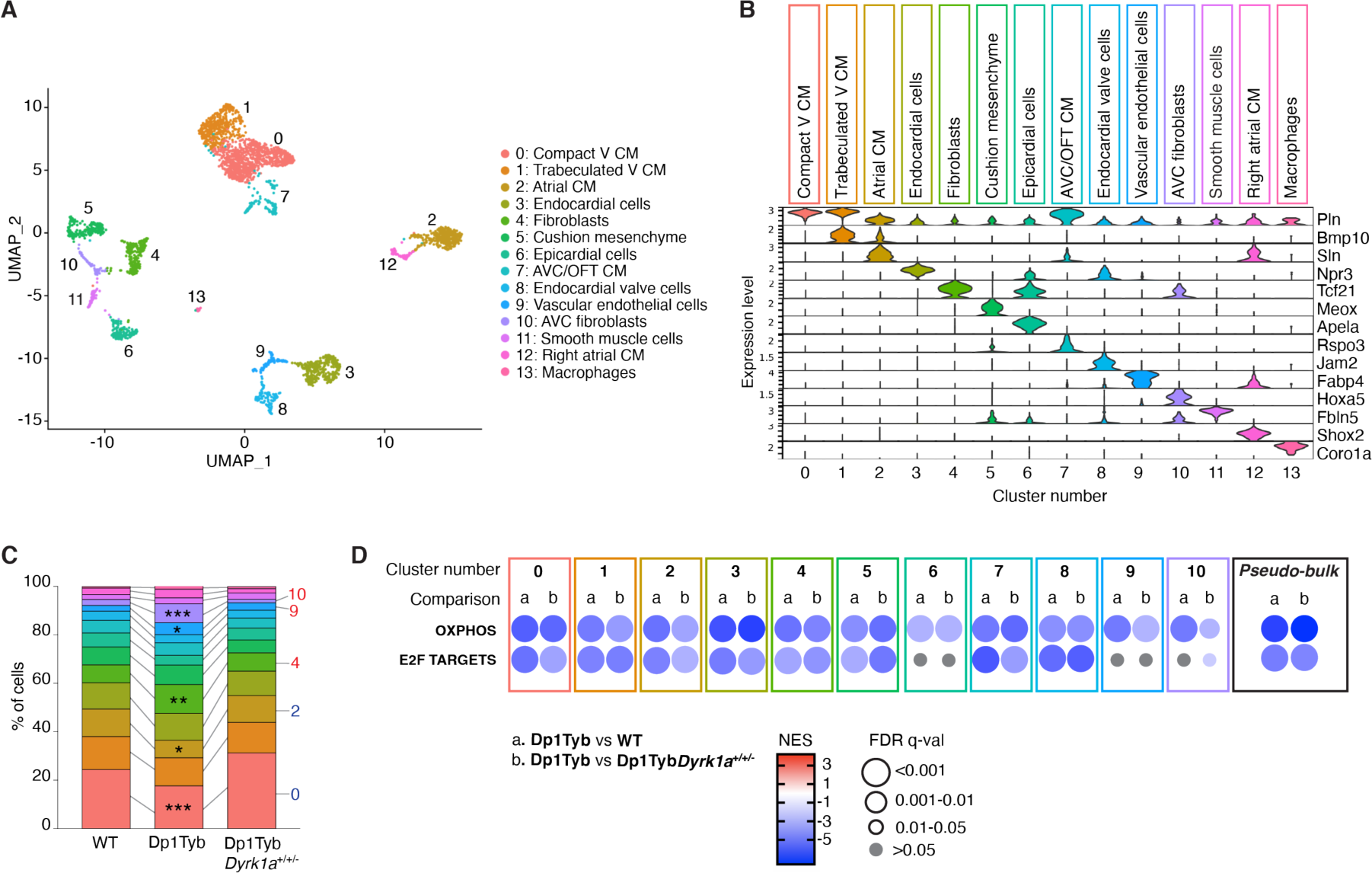
scRNAseq reveals similar gene expression changes across different cell types in Dp1Tyb embryonic hearts. (**A**) Uniform Manifold Approximation and Projection (UMAP) clustering of scRNAseq data pooled from E13.5 hearts WT, Dp1Tyb and Dp1Tyb*Dyrk1a*^+/+/-^ E13.5 hearts. Each dot represents a single cell; colors indicate 14 clusters whose identity was inferred based on expression of markers genes. V, ventricular; CM, cardiomyocytes; AVC, atrioventricular canal; OFT, outflow tract. (**B**) Violin plots showing the expression levels of representative marker genes across the 14 clusters; Y-axis shows the log-scale normalized read count. (**C**) Stacked column plot showing the percentage of cells in each of the 14 cell populations, colored according to cluster designation. Clusters with significantly altered percentages in Dp1Tyb hearts compared to WT are indicated; Fisher’s exact test, * 0.01 < *P* < 0.05, ** 0.001 < *P* < 0.01, *** *P* < 0.001. (**D**) GSEA of Dp1Tyb v WT (a) or Dp1Tyb v Dp1Tyb*Dyrk1a*^+/+/-^ (b) scRNAseq data from the 11 most abundant clusters analyzed individually and pooled (pseudo-bulk), showing key pathways and their NES. Colors and sizes of circles indicate NES and FDR q-value, respectively. Sample numbers: n=2 WT, 1 Dp1Tyb, 2 Dp1Tyb*Dyrk1a*^+/+/-^ embryonic hearts.

To determine which cell types showed the pathway changes seen in whole hearts, we subjected the scRNAseq data to GSEA. Initially, we pooled the scRNAseq across all cell types and found that this pseudo-bulk RNAseq analysis recapitulated the changes seen in the bulk RNAseq analysis, with Dp1Tyb hearts showing decreased expression of oxidative phosphorylation and proliferation genes (Figure 2D). Extending this analysis to the 11 most abundant cell clusters, we found that decreased expression of the oxidative phosphorylation genes was seen in all cell types and decreased expression of proliferation genes in all except epicardial cells, vascular endothelial cells, atrioventricular cushion fibroblasts (Figure 2D). Thus, the key pathway changes are seen in most cell types.

### Proliferative defects in Dp1Tyb embryonic hearts

The decreased cell proliferation signature is present in both human DS and mouse Dp1Tyb hearts. This change is insufficient to cause pathology since it is present in Ts1Rhr hearts which do not have CHD. Nonetheless, it is possible that decreased proliferation contributes to the development of CHD. To investigate directly if cell proliferation was affected in Dp1Tyb embryonic hearts, we measured the fraction of cells in different cell cycle phases. Flow cytometric analysis of Dp1Tyb hearts showed an increased proportion of cardiomyocytes and endocardial cells in G1, and fewer cells in S, consistent with impaired proliferation (Figure 3A-B). This proliferation defect may explain the decreased fraction of cardiomyocytes in Dp1Tyb hearts.

**Figure 3.**
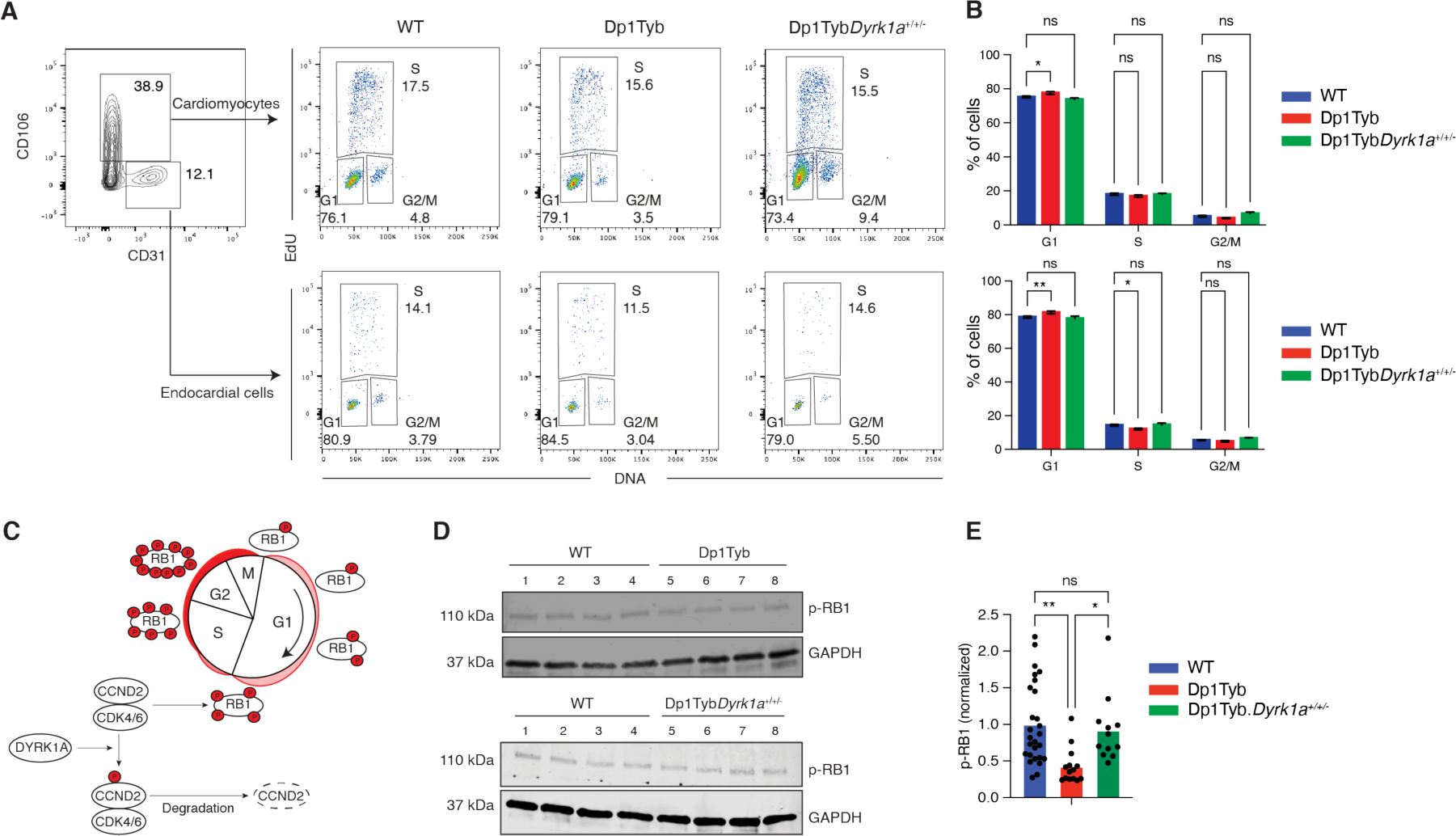
Proliferative defects in Dp1Tyb embryonic hearts. (**A**) Flow cytometric analysis of freshly isolated embryonic hearts pulsed with EdU. EdU and DNA content of cardiomyocytes (CD106^+^CD31^-^) and endocardial cells (CD106^-^ CD31^+^) were used to distinguish cells in G1, S and G2/M phases. Numbers indicate percentage of cells in gates. (**B**) Mean (±SEM) percentage of cells in each cell cycle phase taken from data as in A. (**C**) Diagram showing how DYRK1A may regulate the cell cycle. Cyclin D2 (CCND2) in complex with CDK4/6 phosphorylates RB1 promoting cell cycle progression. DYRK1A phosphorylates CCND2 leading to its degradation thereby causing reduced CDK4/6 activity, reduced RB1 phosphorylation and impaired cell cycle progression. (**D**) Representative immunoblot analysis of lysates from WT, Dp1Tyb and Dp1Tyb.*Dyrk1a^+/+/-^* E13.5 embryonic hearts probed with antibodies to phospho-RB1 (p-RB1) and GAPDH. Each lane represents an individual embryonic heart. (**E**) Mean p-RB1 abundance normalized to GAPDH levels and to the mean of WT samples which was set to 1. Dots represent individual embryos. Statistical significance was calculated using a Kruskal-Wallis test; * 0.01 < *P* < 0.05, ** 0.001 < *P* < 0.01; ns, not significant. Sample sizes: B, n=38 WT, 14 Dp1Tyb, 12 Dp1TybDyrk1a^+/+/-^; E, n=27 WT, 14 Dp1Tyb, 12 Dp1TybDyrk1a^+/+/-^ embryonic hearts.

Pathway analysis of human DS and Dp1Tyb hearts showed downregulated E2F target genes in both species (Figure 1E, G). The E2F transcription factors control the expression of genes required for the G1 to S phase transition of the cell cycle (*12*). Inspection of the genes in the ‘leading edge’ of the E2F target gene set in the GSEA showed that there were 72 and 153 downregulated genes in human DS fetal hearts and Dp1Tyb embryonic hearts respectively, with 49 genes in common between the two species, indicating that downregulation of E2F activity may be the cause of the impaired proliferation (Table S6). The E2F factors are repressed by binding to the Retinoblastoma protein RB1 (*12*). Phosphorylation of RB1 by CDK4 and CDK6 kinases in complex with Cyclin D proteins leads to dissociation of RB1 from E2F proteins allowing the latter to activate genes required for the G1 to S phase transition (Figure 3C). To examine if the reduced E2F activity in Dp1Tyb hearts is due to impaired phosphorylation of RB1, we used immunoblotting to quantitate levels of phosphorylated RB1 (p-RB1). We found that Dp1Tyb hearts had lower levels of p-RB1, which may account for the reduced E2F activity and hence impaired cell proliferation (Figure 3D, E).

### Mitochondrial defects in Dp1Tyb embryonic cardiomyocytes

To determine if the decreased expression of oxidative phosphorylation genes indicated impaired mitochondrial respiration, we used flow cytometry to measure mitochondrial mass and the mitochondrial inner membrane potential. While mitochondrial mass was not altered in either cardiomyocytes or endocardial cells, and mitochondrial potential was unchanged in endocardial cells, the ratio of cells with high to intermediate membrane potential was reduced in cardiomyocytes from E13.5 Dp1Tyb hearts (Figures 4A, B). This decreased potential suggests that the maturation of Dp1Tyb cardiomyocytes may be affected. A developmental transition, typically occurring at E11.5 in the mouse, is characterized by closure of the mitochondrial permeability transition pore, thereby elevating mitochondrial potential, and by elongation and branching of mitochondria (*13*). Confocal microscopy imaging showed that mitochondria in cardiomyocytes from Dp1Tyb E13.5 hearts had lower aspect ratios and form factors compared to those in WT cardiomyocytes, indicating that Dp1Tyb mitochondria are less elongated and branched, consistent with impaired cardiomyocyte maturation (Figure 4C-D, Figure S3A-B). To directly analyze mitochondrial function, we measured mitochondrial respiration of Dp1Tyb embryonic hearts. We found that cells from E13.5 Dp1Tyb hearts had reduced basal and maximal respiration rates, consistent with impaired mitochondrial function (Figures 4E-F). Furthermore, analysis of glycolysis rates showed that cells from Dp1Tyb had increased rates of glycolysis, demonstrating a bioenergetic switch away from oxidative phosphorylation (Figures 4G-H).

**Figure 4.**
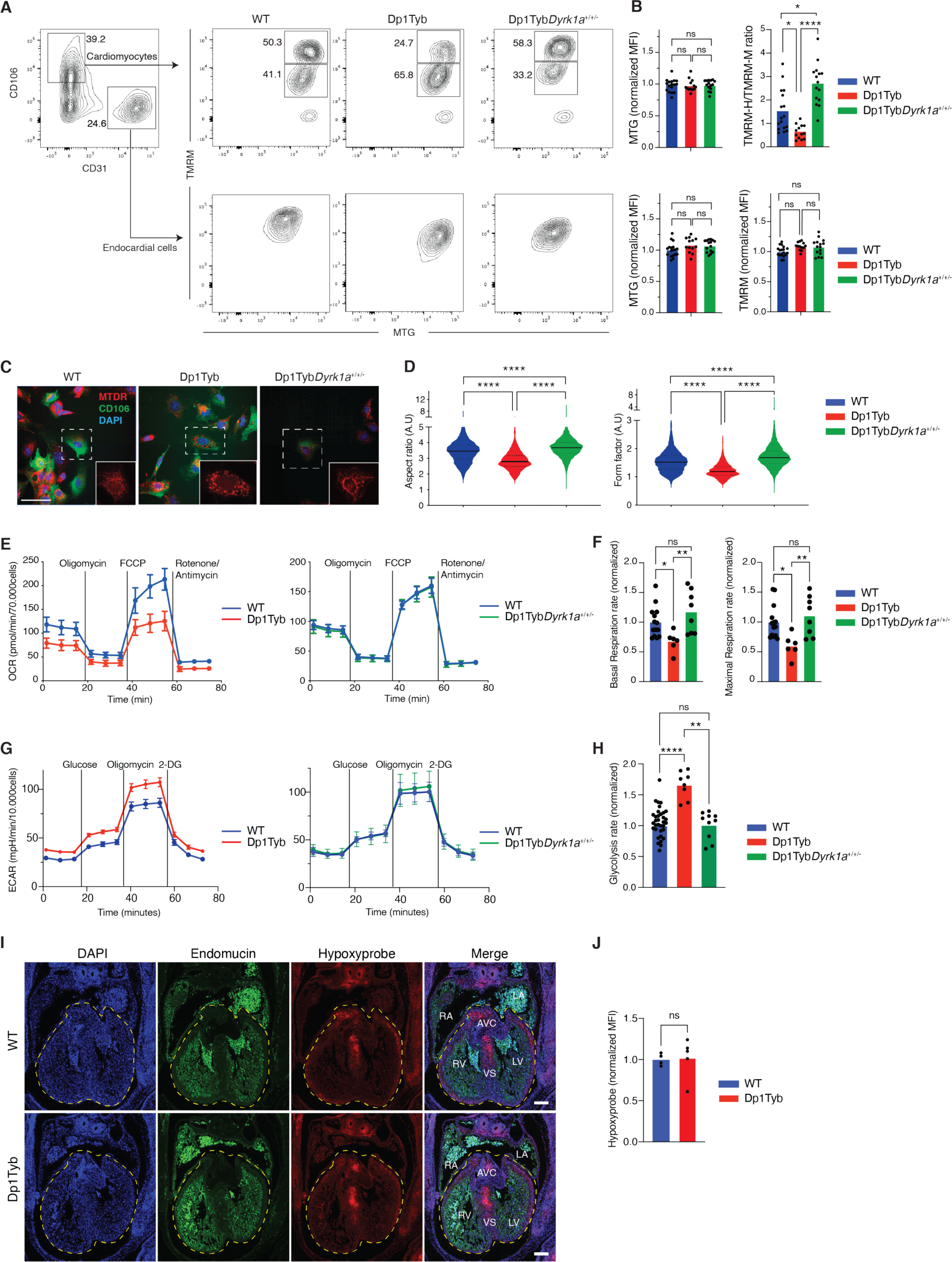
Mitochondrial defects in Dp1Tyb embryonic cardiomyocytes. (**A**) Flow cytometry analysis showing gating strategy used to measure mitochondrial mass (MTG) and mitochondrial potential (TMRM) in cardiomyocytes (CD106^+^CD31^-^) and endocardial cells (CD106^-^CD31^+^) from E13.5 embryonic hearts of the indicated genotypes. Cardiomyocytes were subdivided into cells that have a high (TMRM-H) and medium (TMRM-M) potential. Numbers indicate percentage of cells in gates. (**B**) Mean fluorescence intensity (MFI) of MTG and TMRM in endocardial cells and of MTG in cardiomyocytes normalized to the average of WT samples which was set to 1. For cardiomyocytes mitochondrial potential was measured using a TMRM-H/TMRM-M ratio. Dots represent individual embryos. (**C**) Representative confocal microscopy images of cells from E13.5 hearts of the indicated genotypes showing staining with MitoTracker Deep Red (MTDR - mitochondria), anti-CD106 (cardiomyocytes) and DAPI. Images show a maximum projection of Z-stacks from 0 to 3 µm with a step size of 1 µm. Insets are enlarged images of the region in the dashed square showing the mitochondrial network. Scale bar 50µm. (**D**) Violin plots of mitochondrial aspect ratio and form factor in cardiomyocytes (CD106^+^) determined from images such as those in C (Figures S3A, B). Aspect ratio and form factor are measures of distortion from circularity and degree of branching, respectively (*74*). Black lines indicate median, dotted lines indicate 25th and 75th centiles. (**E**) Mean±SEM oxygen consumption rate (OCR) in E13.5 heart cells from embryos of the indicated genotypes analyzed using a Seahorse analyzer with oligomycin (ATP synthase inhibitor), FCCP (depolarizes mitochondrial membrane potential), and rotenone and antimycin (complex I and III inhibitors) added at the indicated times. Basal respiration rate was calculated from the mean of the first three measurements, maximal respiration rate from the three time points after addition of FCCP. (**F**) Mean basal and maximal respiration rates of E13.5 heart cells normalized to the mean rates in WT hearts. Dots represent individual embryos. (**G**) Mean±SEM extracellular acidification rate (ECAR) in E13.5 heart cells from embryos of the indicated genotypes analyzed using a Seahorse analyzer with glucose, oligomycin (ATP synthase inhibitor) and 2 deoxy-glucose (2-DG, competitive inhibitor of glucose) added at the indicated times. Glycolysis rate was calculated as the difference between the mean ECAR of the three measurements before and after glucose injection. (**H**) Mean glycolysis rates of E13.5 heart cells normalized to the mean rates in WT hearts. Dots represent individual embryos. (**I**) Representative images of sections of E13.5 hearts of the indicated genotypes showing a 4-chamber view stained with anti-Endomucin (endothelial cells), anti-Hypoxyprobe (hypoxia) and DAPI. Dashed line indicates a region of interest (ROI) encompassing the ventricles and the atrioventricular cushions. Scale bar 200µm. (**J**) MFI of anti-Hypoxyprobe in ROI determined from images such as those in I. Dots represent individual embryos. LV, left ventricle; RV, right ventricle; LA, left atrium; RA, right atrium; AVC, atrioventricular cushion; VS, ventricular septum. Statistical significance was calculated using a Kruskal-Wallis (B, D, F, H) or Mann Whitney test (J), * 0.01 < *P* < 0.05, ** 0.001 < *P* < 0.01, **** *P* < 0.0001; ns, not significant. Sample numbers (embryonic hearts): B, n=17 WT, 13 Dp1Tyb, 15 Dp1TybDyrk1a^+/+/-^; D, n=25 WT, 10 Dp1Tyb, 17 Dp1TybDyrk1a^+/+/-^; F, n=14 WT, 6 Dp1Tyb, 8 Dp1Tyb*Dyrk1a*^+/+/-^; H, 35 WT, 8 Dp1Tyb, 10 Dp1TybDyrk1a^+/+/-^; J, n=4 WT, 6 Dp1Tyb.

One possible cause for this switch towards glycolysis could be hypoxia, which through HIF-1α results in the upregulation of genes encoding enzymes in the glycolytic pathway (*14*). To investigate if increased hypoxia could be playing a role in this altered cardiac metabolism, we measured hypoxia in developing Dp1Tyb and control embryonic hearts by injecting pregnant mice with pimonidazole hydrochloride, also known as Hypoxyprobe (*15*). Pimonidazole covalently binds to thiol groups on proteins and amino acids in hypoxic cells and can be visualized with an antibody. This analysis showed no change in the amount of hypoxia in Dp1Tyb hearts (Figure 4I-J). This result was further supported by GSEA which did not show any significant changes in the expression of HIF-1α-regulated genes (Table S6). Thus, increased hypoxia is unlikely to be the cause of the increased glycolysis. Taken together these results demonstrate that Dp1Tyb embryonic hearts have impaired mitochondrial respiration, with a compensatory increase in glycolysis.

### Three copies of *Dyrk1a* required to cause heart defects

To gain further insight into the basis of CHD in DS we used systematic genetic mapping to identify a causative gene responsible for this pathology. Using a panel of mouse strains with a nested series of duplications of regions on Mmu16, we previously showed that a region of 39 genes duplicated in Dp3Tyb was sufficient to cause CHD, but when this was broken down into three shorter duplicated regions in Dp4Tyb, Dp5Tyb and Dp6Tyb mice, none of the resulting strains had heart defects, implying that there must be at least two causative genes (Figure 5A) (*8*). Furthermore, since Ts1Rhr embryos do not have CHD, one of the causative genes must be within the 8 coding genes and 1 microRNA gene that are duplicated in Dp3Tyb but not Ts1Rhr mice (Figure 5A). Another study showed that Dp(16)4Yey mice with an extra copy of a 35-gene region on Mmu16 which partially overlaps Dp3Tyb, also have CHD (*16*). This overlap includes 16 genes at the proximal end of the Dp3Tyb duplication covering all of the Dp4Tyb region and the first two genes within the Dp5Tyb region. Taken together, the simplest explanation for these results is that one causative gene lies within the proximal region duplicated in Dp3Tyb mice but not Ts1Rhr mice, containing two coding genes (*Setd4*, *Cbr1*) and a microRNA gene (*Mir802*) (Figure 5A, orange), and a second causative gene lies within the first two genes of the Dp5Tyb region (*Dyrk1a*, *Kcnj6*) (Figure 5A, blue). Thus, we tested the potential role for each of these 5 genes in causing CHD, by reducing its copy number in Dp1Tyb embryos from three to two.

**Figure 5.**
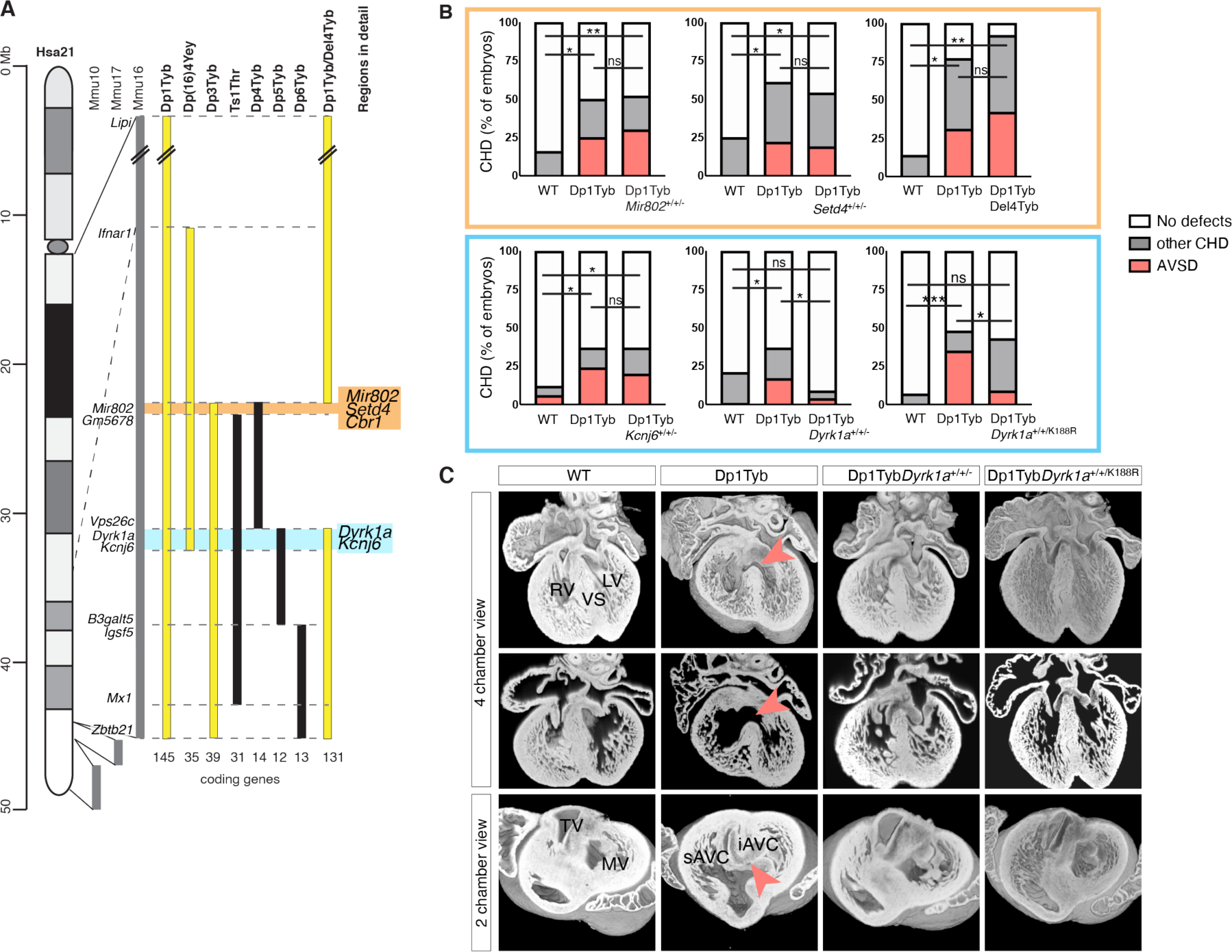
Three copies of *Dyrk1a* required to cause heart defects. (**A**) Map of Hsa21 showing regions of orthology to Mmu10, Mmu17 and Mmu16 (grey) and indicating regions of Mmu16 that are duplicated in mouse strains that show CHD (yellow) and in those that do not (black); genes at boundaries of these duplications are indicated next to the Mmu16 map; numbers of coding genes indicated below duplicated regions. Two genetic intervals containing 3 and 2 candidate genes for CHD are indicated in orange and blue, respectively. (**B**) Graphs of percentage of CHD in E14.5 embryonic hearts from the indicated strains, indicating the frequency of AVSD and other CHD, which are predominantly VSDs and occasionally outflow tract defects such as overriding aorta. n=25 WT, 20 Dp1Tyb, 27 Dp1Tyb*Mir802*^+/+/-^; n=24 WT, 23 Dp1Tyb, 26 Dp1Tyb*Setd4*^+/+/-^; n=7 WT, 13 Dp1Tyb, 12 Dp1TybDel4Tyb; n=32 WT, 41 Dp1Tyb, 30 Dp1Tyb*Kcnj6*^+/+/-^; n=101 WT, 86 Dp1Tyb, 23 Dp1Tyb*Dyrk1a*^+/+/-^; n=30 WT, 23 Dp1Tyb, 44 Dp1Tyb*Dyrk1a*^+/+/K188R^ embryonic hearts. Fisher’s exact test, * 0.01 < *P* < 0.05, ** 0.001 < *P* < 0.01, *** *P* < 0.001 for difference in number of total CHD, except for Dp1Tyb*Dyrk1a*^+/+/K188R^ cohort, where statistics were calculated for number of AVSD; ns, not significant. (**C**) 3D HREM rendering of WT, Dp1Tyb, Dp1Tyb*Dyrk1a*^+/+/-^ and Dp1Tyb*Dyrk1a*^+/+/K188R^ hearts, eroded to show an anterior four-chamber view (top and middle) and a two-chamber view (bottom) seen from the atria at the level of the atrioventricular canal. Top and bottom rows show eroded 3D views, middle row shows 2D sections at the same level as shown in the top row. Red arrowheads indicate VSD (top, middle) or AVSD (bottom). iAVC, inferior atrio-ventricular cushion; LV, left ventricle; MV, mitral valve; RV, right ventricle; sAVC, superior atrio-ventricular cushion; TV, tricuspid valve; VS, ventricular septum.

We crossed Dp1Tyb mice to mouse strains deficient in each of the candidate genes, except for *Cbr1*, which we tested by crossing to Del4Tyb mice that have a deletion of the entire Dp4Tyb region (Figure 5A, Figure S4A). For each cross we analyzed the hearts of E14.5 embryos using high resolution episcopic microscopy (HREM) (*17*) to generate detailed 3D images. The use of such a 3D technique is essential to accurately identify different types of CHD. As expected, compared to WT controls, Dp1Tyb embryos had significantly increased rates of CHD, especially the more severe AVSD, which are typified by a common atrioventricular valve in place of separated mitral and tricuspid valves (Figure 5B-C). Removing one copy of *Mir802*, *Setd4*, *Cbr1* or *Kcnj6* did not affect the frequency of CHD in general, or specifically AVSDs. However, reducing the copy number of *Dyrk1a* from three to two completely rescued CHD in Dp1Tyb*Dyrk1a*^+/+/-^ mice (Figures 5B-C). Thus, three copies of *Dyrk1a* are required to cause CHD in Dp1Tyb mice.

*Dyrk1a* encodes the DYRK1A serine/threonine protein kinase. To investigate whether increased DYRK1A kinase activity is required for CHD, we crossed Dp1Tyb mice to a strain carrying an allele coding for kinase-inactive DYRK1A (*Dyrk1a*^K188R^). The resulting Dp1Tyb*Dyrk1a*^+/+/K188R^ embryos had no increase in AVSDs compared to WT controls, and significantly fewer AVSDs than Dp1Tyb embryos (Figures 5B-C). Thus, increased DYRK1A catalytic kinase activity is required for severe CHD in Dp1Tyb mice.

### Increased dosage of *Dyrk1a* causes key transcriptional changes in Dp1Tyb embryonic hearts

Next, we investigated if increased dosage of *Dyrk1a* is responsible for the transcriptional changes in Dp1Tyb E13.5 embryonic hearts. A comparison of the transcriptomes of WT, Dp1Tyb and Dp1Tyb*Dyrk1a*^+/+/-^ hearts showed that compared to WT controls, Dp1Tyb*Dyrk1a*^+/+/-^ samples had increased expression of the duplicated genes, but very few other differentially expressed genes (Figure 6A, Table S7). In contrast, comparison of Dp1Tyb and Dp1Tyb*Dyrk1a*^+/+/-^ hearts showed large numbers of differentially expressed genes, as had been seen in the comparison of Dp1Tyb with WT controls. These results show that an extra copy of *Dyrk1a* is responsible for most of the transcriptional changes in developing Dp1Tyb hearts. We extended this analysis to the proteome using mass spectrometric analysis of Dp1Tyb and Dp1Tyb*Dyrk1a*^+/+/-^ E13.5 hearts, revealing 292 differentially expressed proteins (Figure 6B, Table S8). As expected, the level of DYRK1A was about 1.5-fold higher in Dp1Tyb compared to Dp1Tyb*Dyrk1a*^+/+/-^ hearts. Moreover, pathway analysis of the transcriptomic and proteomic data showed that 3 copies of *Dyrk1a* are required for the decreased expression of oxidative phosphorylation and proliferation genes and increased expression of EMT genes, but not for the increased expression of immune response genes (Figure 6C, Figure S1D, E). A comparison of the differentially expressed genes in Dp1Tyb v WT hearts and Dp1Tyb v Dp1Tyb*Dyrk1a*^+/+/-^ hearts, showed that there were 34 upregulated and 47 downregulated genes in common between these comparisons, representing genes most likely to be regulated by increased *Dyrk1a* dosage (Table S9). STRING protein interaction (https://string-db.org/) analysis of these genes once again showed a clear signature of decreased networks of proteins involved in proliferation.

**Figure 6.**
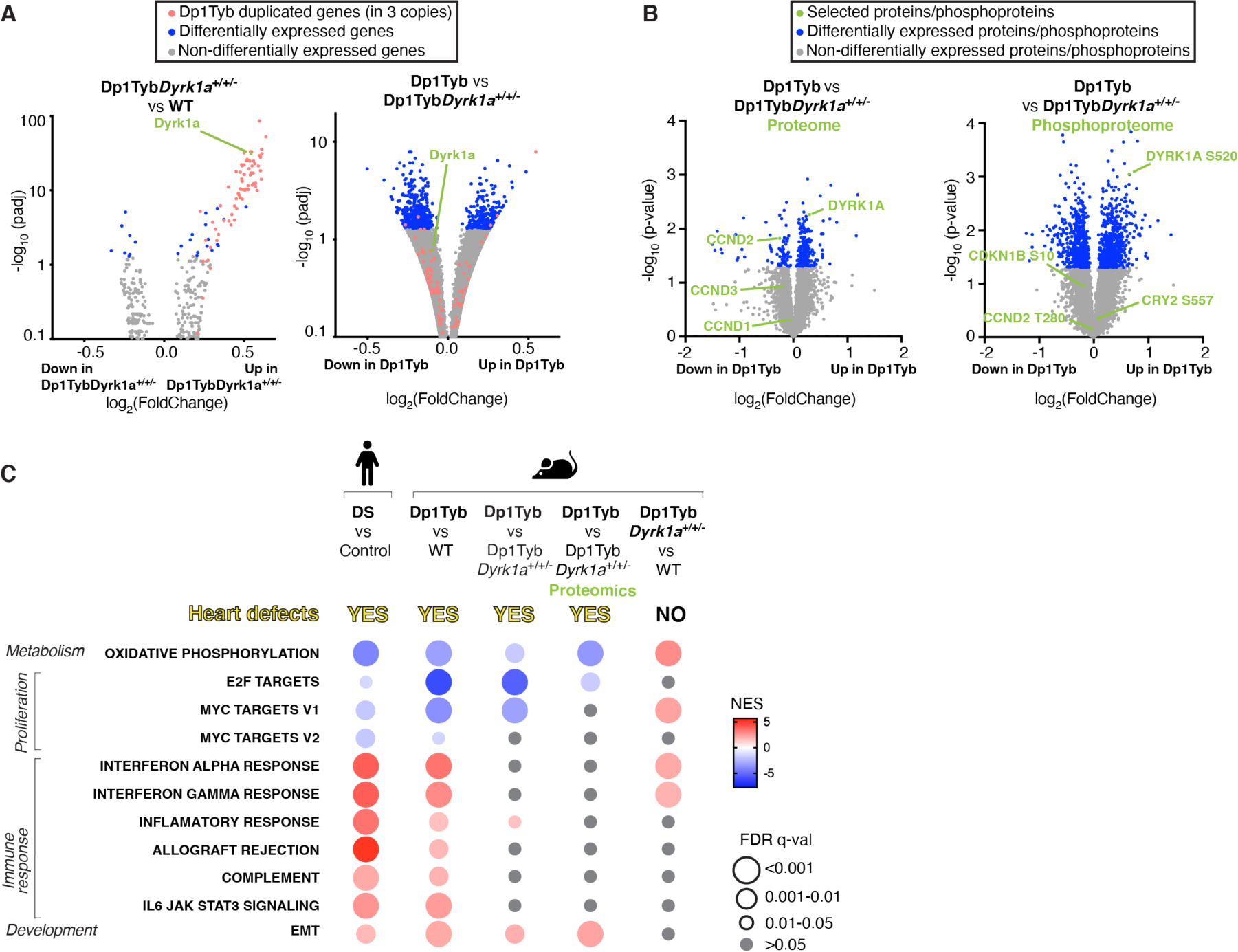
Increased dosage of *Dyrk1a* causes key transcriptional changes in Dp1Tyb embryonic hearts. (**A**) Volcano plots showing fold-change in gene expression in E13.5 mouse embryonic hearts (Dp1Tyb*Dyrk1a*^+/+/-^ v WT and Dp1Tyb v Dp1Tyb*Dyrk1a*^+/+/-^) plotted against adjusted *P*-value for significance of the difference. Genes present in three copies in Dp1Tyb mice (red) and differentially expressed genes (blue) and *Dyrk1a* (green) are indicated. (**B**) Volcano plots showing fold-change in abundance of proteins and phosphorylated sites in Dp1Tyb v Dp1Tyb*Dyrk1a*^+/+/-^ E13.5 hearts. DYRK1A, CCND1, CCND2 and CCND3 are indicated in green on the proteome plot; phosphorylated sites known to be DYRK1A targets and an autophosphorylation site on DYRK1A are indicated on the phosphoproteome plot. (**C**) Comparison of dysregulated pathways determined by GSEA of RNAseq and proteomic experiments. Colors and sizes of circles indicate NES and FDR q-value, respectively. Sample numbers: n=5 embryonic hearts.

Furthermore, scRNAseq analysis of Dp1Tyb*Dyrk1a*^+/+/-^ E13.5 hearts showed that reduction of the copy number of *Dyrk1a* reversed the decreased fraction of cardiomyocytes and the increased fraction of fibroblasts seen in Dp1Tyb hearts (Figure 2C, Figure S2), demonstrating that these changes in cell type abundance are dependent on three copies of *Dyrk1a*. Moreover, GSEA pathway analysis showed that changes in expression of oxidative phosphorylation, proliferation and EMT genes which are seen across most cell types, are also dependent on 3 copies *Dyrk1a* (Figure 2D).

### Broad expression of *Dyrk1a* in many cardiac cell types

Since a third copy of *Dyrk1a* has a profound effect on the transcriptomes of most cell types in the developing heart, we examined the expression pattern of this key gene. Using the scRNAseq data, we found that *Dyrk1a* is expressed in all cell types in E13.5 hearts, a conclusion that was further supported by RNAscope analysis of E12.5 and E14.5 hearts, an *in situ* hybridization method that detects mRNA in tissue sections (Figure 7A-B). This broad expression of *Dyrk1a* is consistent with a role for increased DYRK1A causing the observed pathway changes across multiple cell types.

**Figure 7.**
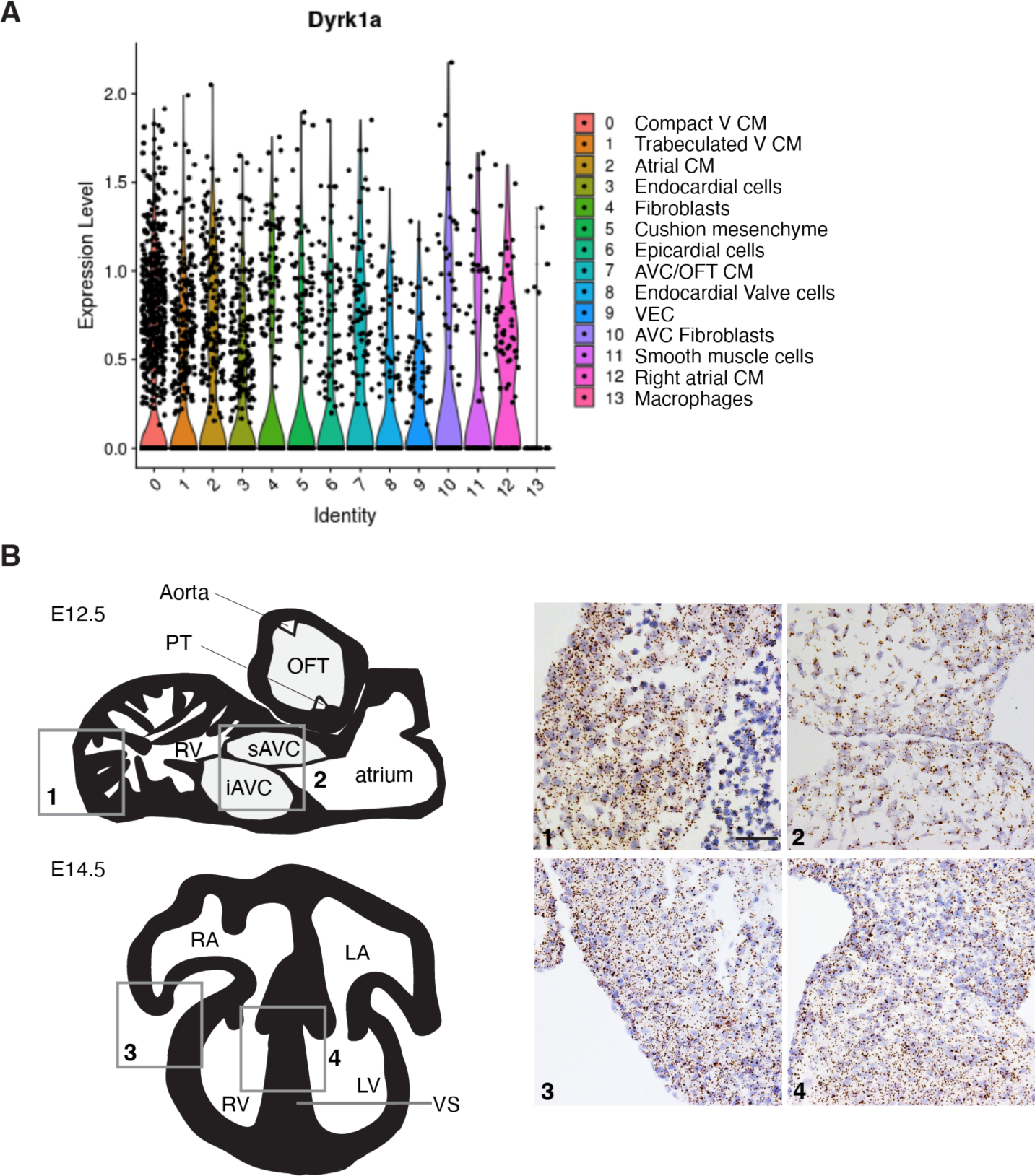
*Dyrk1a* expression in a broad range of cell types in the developing heart. (**A**) Violin plots showing expression of *Dyrk1a* in mouse E13.5 hearts across 14 cell clusters identified in Figure 2B. Dots indicate single cells. Sample numbers: n=5 embryonic hearts. (**B**) Left, schematic illustrations of a sagittal section of an E12.5 heart and a 4-chamber view of an E14.5 heart. Right, RNAscope analysis of *Dyrk1a* expression (brown dots) in sections of the right ventricle myocardial wall (1) and atrioventricular cushions (2) at E12.5 and right ventricular myocardial wall (3) and ventricular septum (4) at E14.5. *Dyrk1a* expression is ubiquitous. Scale bar 50 µm. AV, atrioventricular; iAVC, inferior atrioventricular cushion; LA, left atrium; LV, left ventricle; OFT, outflow tract; PT, pulmonary trunk; RA, right atrium; RV, right ventricle; sAVC, superior atrioventricular cushion; VS, ventricular septum.

### Increased dosage of *Dyrk1a* is required for reduced proliferation in Dp1Tyb embryonic hearts

Next, we investigated if increased dosage of *Dyrk1a* was also required for the physiological changes seen in Dp1Tyb embryonic hearts. Analysis of proliferation showed that Dp1Tyb*Dyrk1a*^+/+/-^ cardiomyocytes and endocardial cells showed no change in the fraction of cells in G1, S and G2/M compared to WT cells, demonstrating that 3 copies of *Dyrk1a* are required for the impaired proliferation (Figures 3A, B). Furthermore, the reduction in phosphorylated RB1 in Dp1Tyb embryonic hearts was also reversed by reducing the copy number of *Dyrk1a* from 3 to 2 (Figure 3D-E), and Dp1Tyb*Dyrk1a*^+/+/-^ hearts no longer showed reduced expression of E2F target genes (Figure 6C, Table S6). Thus, a third copy of *Dyrk1a* is required for the reduced phosphorylation of RB1 and the decreased expression of E2F target genes in Dp1Tyb hearts.

Increased expression of DYRK1A has been previously suggested to cause decreased cell proliferation by phosphorylating Cyclin D (CCND) proteins, leading to their degradation (Figure 3C) (*18–22*). To investigate if this might be occurring in Dp1Tyb hearts we used mass spectrometry to analyze the phospho-proteome of the mutant hearts. Comparison of Dp1Tyb and Dp1Tyb*Dyrk1a*^+/+/-^ E13.5 hearts showed that there were many phospho-peptides whose abundance differed significantly because of a third copy of *Dyrk1a*, including increased pS520-DYRK1A, an autophosphorylation site of the kinase, consistent with increased DYRK1A activity in Dp1Tyb compared to Dp1Tyb*Dyrk1a*^+/+/-^ hearts (*23*). However, we saw no significant change in the abundance of pT280-CCND2, a site reported to be phosphorylated by DYRK1A in cardiomyocytes (Figure 6B, Table S8) (*20*). Furthermore, we were able to detect only two other known DYRK1A phosphorylation targets (p-S10-CDKN1B and p-S557-CRY2), neither of which was significantly altered in abundance. However, analysis of the whole proteome showed significantly decreased levels of CCND2 in Dp1Tyb hearts, and decreased levels of CCND1 and CCND3, albeit these latter changes were not significant (Figure 6B). We speculate that increased DYRK1A causes elevated phosphorylation of CCND2 leading to its rapid degradation, explaining the lack of change in the abundance of phosphorylated CCND2, while resulting in reduced amounts of CCND2. These results are consistent with increased dosage of DYRK1A impairing proliferation in Dp1Tyb embryonic hearts through phosphorylation and subsequent degradation of CCND2, leading to reduced CDK4/6-induced phosphorylation of RB1 and hence less expression of E2F-regulated genes that are required for G1 to S phase transition.

### Three copies of *Dyrk1a* cause mitochondrial dysfunction in embryonic cardiomyocytes

Next, we examined if the mitochondrial dysfunction in Dp1Tyb cardiomyocytes was dependent on 3 copies of *Dyrk1a*. We found that reduction of the copy number of *Dyrk1a* from 3 to 2 reversed the decreased mitochondrial potential and impaired mitochondrial morphology in Dp1Tyb cardiomyocytes (Figures 4A-D). Furthermore, cells from Dp1Tyb*Dyrk1a*^+/+/-^ E13.5 hearts showed normal rates of basal and maximal respiration and glycolysis (Figures 4E-H). Taken together, these results show that increased expression of *Dyrk1a* causes impaired mitochondrial function in Dp1Tyb embryonic cardiomyocytes.

### Pharmacological inhibition of DYRK1A results in a partial rescue of heart defects in Dp1Tyb embryos

Finally, we investigated if treatment of pregnant mice with an inhibitor of DYRK1A kinase activity could reverse the CHD in Dp1Tyb embryos. We used Leucettinib-21, a recently developed DYRK1A inhibitor and a negative control, iso-Leucettinib-21, an inactive isomer (Figure S5A). Leucettinib-21 is a potent inhibitor of DYRK1A (IC_50_ = 3.1 nM), whereas iso-Leucettinib-21 has an IC_50_ >10 µM for DYRK1A (*24*). Leucettinib-21 shows strong selectivity for DYRK1A and related kinases. In a screen of 413 human kinases, only DYRK1A and 5 other kinases were inhibited by Leucettinib-21 with IC_50_ values of <10 nM (L. Meijer, personal communication). First, we asked if Leucettinib-21 administered to the pregnant mouse could pass through the placenta into the embryo. Analysis of mice treated with 0.3, 3 or 30 mg/kg of Leucettinib-21 showed detectable Leucettinib-21 in the embryo at 3 and 30 but not 0.3 mg/kg (Figure S5B). Based on the measured amounts of Leucettinib-21 in the embryo, the 3 and 30 mg/kg doses result in around 24 nM and 560 nM Leucettinib-21 in the embryo. Since the IC_50_ for inhibiting DYRK1A in cultured cells is estimated to be around 36 nM (L. Meijer, personal communication), we chose to use 30 mg/kg Leucettinib-21 to achieve a concentration of Leucettinib-21 in the embryo above this cellular IC_50_ value.

We treated pregnant mice with Leucettinib-21 or iso-Leucettinib-21 daily starting at E5.5 which is before the heart forms until E13.5 for RNAseq analysis and E14.5 for analysis of CHD by HREM (Figure 8A). Homozygous genetic deletion of *Dyrk1a* results in a mid-gestation lethality, with no embryos surviving beyond E13.5 (*25*). Thus, it was possible that Leucettinib-21 treatment would cause embryonic lethality. However, we found that treatment of mated mice with 30 mg/kg Leucettinib-21 did not affect the fraction of mice that became pregnant or the size of their litters (Figure S5C-D). Furthermore, the recovery of Dp1Tyb embryos at E14.5 was also not affected (Figure S5E), in line with the normal segregation of the mutation, as previously reported (*8*). Thus, Leucettinib-21 treatment does not interfere with pregnancy and is likely only partially inhibiting DYRK1A.

**Figure 8.**
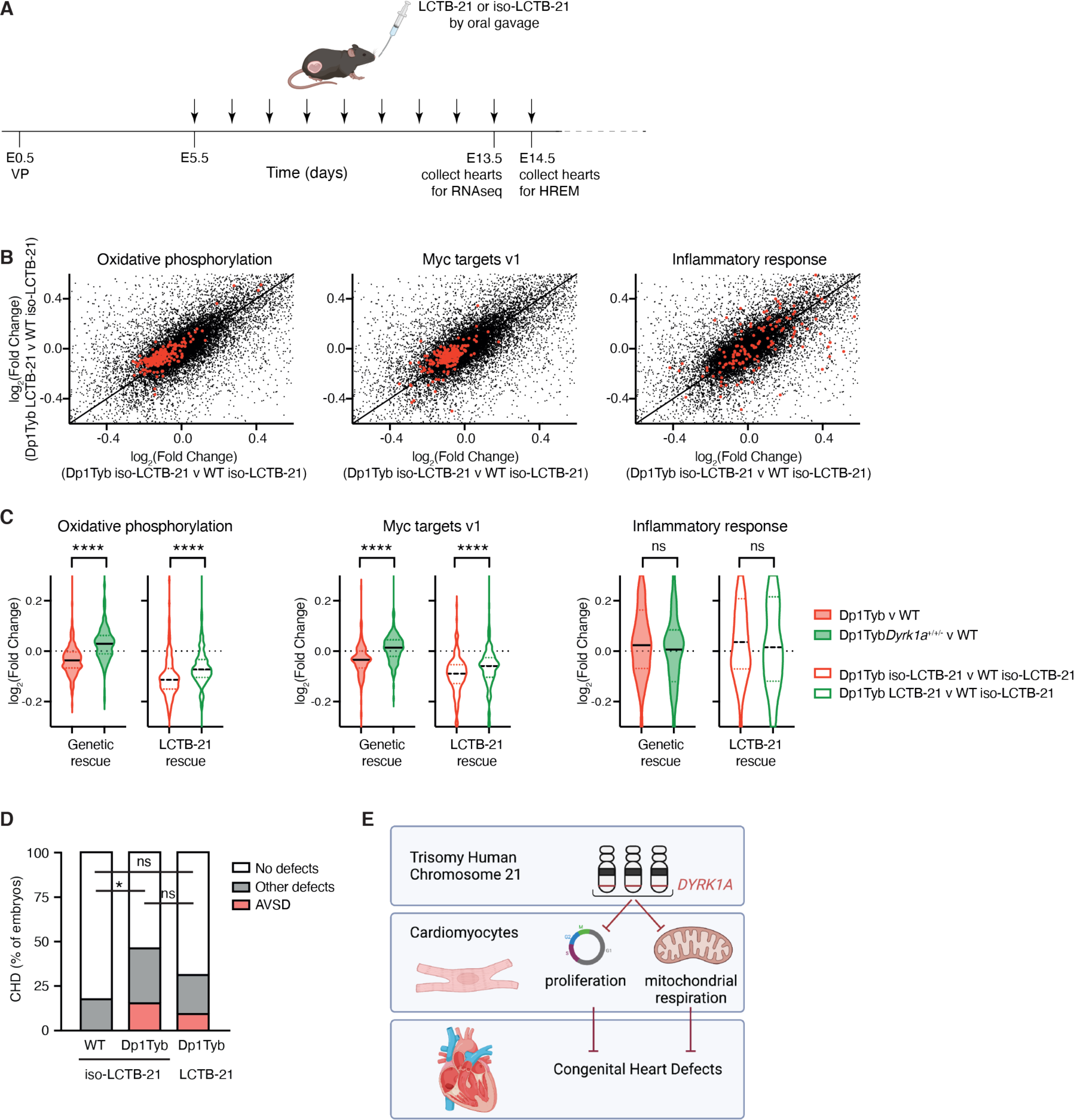
Pharmacological inhibition of Dyrk1a partially rescues CHD in the Dp1Tyb mouse model of DS. (**A**) C57BL/6J females that had been mated with Dp1Tyb males, were treated daily by oral gavage with Leucettinib-21 (LCTB-21) or iso-Leucettinib-21 (iso-LCTB-21) from 5 days after vaginal plug was found (embryonic day 0.5, E0.5). Embryos were collected at E13.5 for RNAseq, or at E14.5 for HREM. (**B**) Scatter plots comparing the log_2_(fold-change) (Log2FC) of mRNA levels for all genes (black) for Dp1Tyb v WT embryos both treated with iso-LCTB-21 or Dp1Tyb embryos treated with LCTB-21 v WT embryos treated with iso-LCTB-21. Genes from the Hallmark genesets for Oxidative Phosphorylation, Myc targets V1 and inflammatory response, are highlighted in red. (**C**) Violin plots showing the Log2FC of expression of the same genesets as in B, for the following comparisons: Dp1Tyb v WT, Dp1TybDyrk1a^+/+/-^ v WT, Dp1Tyb treated with iso-LCTB-21 v WT treated with iso-LCTB-21 and Dp1Tyb treated with LCTB-21 v WT treated with iso-LCTB-21. Black lines indicate median, dotted lines indicate 25th and 75th centiles. Dotted line at 0 indicates no change. (**D**) Graph of percentage of CHD in E14.5 embryonic hearts from the indicated strains. (**E**) Three copies of *DYRK1A* lead to impaired proliferation and mitochondrial respiration in cardiomyocytes and congenital heart defects. Created in Biorender. Statistical tests were carried out with a Kruskal-Wallis (C) or Fisher’s exact (D) test; * 0.01 < *P* < 0.05, **** *P* < 0.0001; ns, not significant. Sample numbers (embryos): B, C, n=5 for each condition; D, n=34 WT, 26 Dp1Tyb treated with iso-LCTB-21, 32 Dp1Tyb treated with LCTB-21.

RNAseq analysis of E13.5 embryos from mice treated with Leucettinib-21 or iso-Leucettinib-21 showed that the active Leucettinib-21 inhibitor partially reversed the decreased expression of genes in the oxidative phosphorylation and proliferation (Myc targets v1) pathways in Dp1Tyb embryonic hearts, however, it had no significant effect on the increased expression of inflammatory genes in the mutant hearts (Figure 8B, Table S10). The changes in expression caused by Leucettinib-21 were qualitatively similar to the changes seen when the copy number of *Dyrk1a* was reduced from 3 to 2 in Dp1Tyb embryos but were smaller in magnitude, consistent with Leucettinib-21 causing a partial inhibition of DYRK1A in the embryonic heart (Figure 8C). Lastly, we used HREM to analyze the embryos for CHD. We found that Leucettinib-21 treatment resulted in a partial reduction in the frequency of CHD in Dp1Tyb embryos, again consistent with a partial inhibition of DYRK1A (Figure 8D). Thus, pharmacological treatment of pregnant mice with Leucettinib-21 results in a partial rescue of the DYRK1A-dependent CHD in Dp1Tyb embryos.

## Discussion

Our results show that DS fetal hearts have characteristic transcriptional changes, many of which are shared with embryonic hearts from the Dp1Tyb and Dp3Tyb mouse models of DS that show CHD. Gene set enrichment analysis identified several pathways that are altered in the hearts of both human DS and mouse models. Of these, decreased expression of oxidative phosphorylation genes correlated most strongly with CHD in the mouse models, suggesting that impaired mitochondrial function may be an important cause of the developmental defects. In agreement with our results, decreased expression of oxidative phosphorylation genes in DS fetal hearts has been previously seen using microarray technology (*26*). Importantly, physiological analysis demonstrated reduced mitochondrial membrane potential and respiration in Dp1Tyb embryonic cardiomyocytes. Decreased expression of proliferation genes partially correlates with CHD, being seen in human DS hearts, and mouse Dp1Tyb, Dp3Tyb hearts. Hearts from the Ts1Rhr strain which does not have CHD also showed decreased expression of proliferation genes, but only in two out of three gene sets compared to a decrease in all three in Dp1Tyb and Dp3Tyb hearts, suggesting that impaired proliferation may also play an important role in CHD. Changes in several other pathways -increased expression of inflammatory, interferon response and EMT genes - were also seen in the human and mouse embryonic hearts. Since changes in these pathways were seen in hearts from Ts1Rhr mice, they are not sufficient on their own to cause heart defects. However, they may also contribute to CHD pathology, in combination with the mitochondrial and proliferative deficits.

It remains unclear how defects in mitochondrial function and proliferation in most cardiac cells can cause localized defects in septation, rather than a broader cardiomyopathy. One possibility is that the cellular changes are relatively small and that they preferentially affect structures involved in septation such as the ventricular septum, the atrioventricular and outflow tract cushions or the dorsal mesenchymal protrusion. Further work will be needed to understand the developmental defects in Dp1Tyb hearts that lead to VSD and AVSD.

Mitochondrial dysfunction has been previously reported in DS (*27–29*). Human DS fibroblasts, astrocytes and neurons have impaired mitochondrial respiratory activity, decreased mitochondrial membrane potential and altered mitochondrial shape with smaller and more fragmented mitochondria (*30–33*). This dysfunction may underlie the neurological and cognitive impairment in DS, contributing to decreased neurogenesis and altered processing of APP, leading to deposition of Aβ amyloid and early-onset Alzheimer’s disease. Our results show that a very similar mitochondrial phenotype is evident in embryonic cardiomyocytes from the Dp1Tyb mouse model of DS and suggest that mitochondrial dysfunction may play an important role in the cardiac pathology in DS. This commonality of mitochondrial impairment across multiple DS tissues has led to the interesting proposal that drugs that increase mitochondrial biogenesis or respiratory capacity may be promising therapeutic candidates for DS phenotypes, for example, for cognitive deficits (*27–29*). Our results suggest that a similar approach ameliorate cardiac defects.

We had previously shown that Dp3Tyb embryos have CHD, but Dp4Tyb, Dp5Tyb and Dp6Tyb embryos do not, implying that there must be at least two causative genes whose increased dosage leads to CHD (*8*). We now demonstrate that crossing Del4Tyb to Dp1Tyb does not affect the frequency of CHD, thus three copies of the genes in the Dp4Tyb region are not required for cardiac defects and the causative genes must lie in the regions duplicated in Dp5Tyb and Dp6Tyb mice, regions A and B respectively, with at least one causative gene in each region (the 2-locus hypothesis, Figure S6). Using systematic genetic mapping, we now show that one of the causative genes in region A is *Dyrk1a*, although there may be other causative genes in this region. The second causative gene, lying in region B, is likely to be one of the six coding genes present in three copies in Dp3Tyb and Dp6Tyb mice but not Ts1Rhr mice (*Mx2*, *Tmprss2*, *Ripk4*, *Prdm15*, *C2cd2* and *Zbtb21*). None of these six genes has been previously implicated in the etiology of CHD. Interestingly, we note that a recent study of rare copy number variants associated with AVSD identified several patients with and extra copy of regions of Hsa21 ranging from 10-21 Mb and spanning both regions that we have identified in this mouse study, and proposed *DYRK1A* as a candidate gene for CHD (*34*).

Our data show that increased dosage of *Dyrk1a* is required but not sufficient to cause the mitochondrial defects and CHD, since these are not seen in Ts1Rhr embryonic hearts despite 3 copies of *Dyrk1a*. It is unclear how elevated DYRK1A activity, acting with the unknown other causative gene(s), causes mitochondrial changes and CHD. One possibility is that increased DYRK1A inhibits the function of NFAT transcription factors, since DYRK1A phosphorylates NFAT proteins leading to their nuclear exclusion (*35*), and mouse embryos deficient in both NFATc3 and NFATc4 have impaired mitochondrial function and defective cardiac development (*36*). Indeed, DYRK1A in collaboration with RCAN1, another Hsa21-encoded gene, negatively regulates NFAT proteins and overexpression of both genes in the developing mouse leads to failure of outflow tract valve elongation (*35*). While these are not the same defects as seen in DS, which are usually defects in septation, it supports the view that DYRK1A, acting through NFAT proteins can perturb cardiac development. Alternatively, DYRK1A may affect mitochondrial function through the SIRT1 protein deacetylase, since DYRK1A binds to, phosphorylates and activates SIRT1 and overexpression of SIRT1 in cardiomyocytes impairs their mitochondrial respiration (*37, 38*). SIRT1 in turn may be acting through PGC1α (PPARGC1A), a regulator of mitochondrial biogenesis (*39, 40*).

Increased *Dyrk1a* dosage is also required for the impaired proliferation, potentially by phosphorylating Cyclin D proteins leading to their degradation. This in turn would result in reduced CDK4/6 activity, decreased phosphorylation of RB1, reduced activity of E2F transcription factors and less expression of E2F target genes which are required for G1 to S phase transition. The experimental results presented here support such a mechanism.

Furthermore, our conclusions are in agreement with previous studies. Overexpression of DYRK1A in the cardiomyocytes of adult mice leads to reduced levels of CCND2, RB1 phosphorylation and RB1/E2F signaling and hence to impaired proliferation (*20*). Conversely, inhibition of DYRK1A or cardiomyocyte-specific deletion of the *Dyrk1a* gene leads to increased cardiomyocyte proliferation and cardiac hypertrophy in adult mice (*41*). DYRK1A has been shown to phosphorylate LIN52, leading to assembly of the repressive form of the DREAM complex which promotes entry into quiescence, providing another mechanism by which DYRK1A overexpression may regulate proliferation (*42, 43*).

DYRK1A phosphorylates many proteins (*44, 45*); for example, in addition to the substrates described above (NFAT, SIRT1, CCND2 and LIN52), DYRK1A phosphorylates the C-terminal domain of RNA polymerase II, Alternative Splicing Factor (ASF) and CAS9, thereby regulating transcription, splicing and apoptosis (*46–49*). Further studies will be needed to determine whether any of these DYRK1A targets is involved in DYRK1A-induced CHD.

Three copies of *Dyrk1a* are not required for the increased expression of inflammatory and interferon response genes, since these changes were still seen in Dp1Tyb*Dyrk1a*^+/+/-^ hearts. It has been proposed that the increased expression of interferon response genes in DS is caused by a third copy of four interferon receptor genes located on Hsa21 (*50*). Since these expression changes are also seen in Dp3Tyb and Ts1Rhr hearts which do not have an extra copy of the interferon receptor genes, our results show that one or more of the 31 genes present in three copies in Ts1Rhr mice can also cause these inflammatory gene expression changes. A recent report has shown that a third copy of the interferon receptor genes causes CHD in DS (*51*). However, since Dp3Tyb mice that do not have an increased dosage of these genes show CHD (*8*), and Dp1Tyb*Dyrk1a*^+/+/-^ mice that still have three copies of the genes do not show CHD (this study), our results imply that increased dosage of the interferon receptor genes is neither required nor sufficient to cause CHD. Nonetheless it is possible that cardiac pathology results from a complex genetic interplay between *Dyrk1a*, the interferon receptor genes and other unknown causative genes.

To investigate the translational potential of *Dyrk1a* as a causative gene for CHD, we tested whether a pharmacological inhibitor of DYRK1A could reverse the transcriptional changes and heart defects in Dp1Tyb embryonic hearts. Treatment of pregnant mice with Leucettinib-21 resulted in a partial reversal of the transcriptional changes in Dp1Tyb embryos, with effects limited to pathways whose changes were dependent on an extra copy of *Dyrk1a*. Leucettinib-21 treatment also resulted in a small reduction in the frequency of CHD in Dp1Tyb embryos, consistent with a partial inhibition of DYRK1A in the embryos. Further work is needed to understand why the inhibitor does not have a stronger effect, but possibilities include limited access of Leucettinib-21 to embryonic cardiac cells and pharmacokinetic issues such as a short half-life of the inhibitor in the embryo. These could potentially be addressed with higher doses of the inhibitor or more frequent dosing.

In conclusion, our results have, for the first time, revealed genetic and cellular mechanisms leading to CHD in DS. We show that three copies of *Dyrk1a* and increased activity of DYRK1A kinase are required for CHD and result in impaired cardiomyocyte proliferation and mitochondrial respiration. Thus, we propose that CHD in DS arise in part from increased DYRK1A activity in cardiomyocytes leading to reduced proliferation and mitochondrial dysfunction (Figure 8E). Importantly, our work has revealed a therapeutic target for this common human pathology. Furthermore, our systematic genetic mapping approach for dosage-sensitive genes is a paradigm that can be used to identify causative genes and mechanisms responsible for the many other phenotypes of DS.

## Materials and Methods Mice

Mice carrying the following alleles have been described previously: Dp(16Lipi-Zbtb21)1TybEmcf (Dp1Tyb), Dp(16Mir802-Zbtb21)3TybEmcf (Dp3Tyb), Dp(16Cbr1-Fam3b)1Rhr (Ts1Rhr), *Kcnj6*^tm1Stf^ (*Kcnj6*^-^), and *Dyrk1a*^tm1Mla^ (*Dyrk1a*^-^) (*8, 25, 52, 53*). Dp1Tyb mice have been deposited with JAX (strain #037183). Mice with the Del(16Mir802-Vps26c)4TybEmcf (Del4Tyb) allele were generated using an *in vivo* Cre-mediated recombination strategy by breeding female mice containing the *Hprt*^tm1(cre)Mnn^ allele (*54*) and two loxP sites located in trans configuration on Mmu16 at the boundaries of the desired deletion, to C57BL/6J males. Cre activity in the female germline from the *Hprt*^tm1(cre)Mnn^ allele resulted in occasional pups (5.3%) with recombination between the loxP sites generating the Del4Tyb deletion. Mice carrying the loxP sites were derived from targeting ES cells with MICER vectors MHPN219l02 (16:92850909 – 16:92859345 Mb, coordinates from mouse genome assembly GRCm39), located between *Runx1* and *Mir802,* and MHPP432c09 (16:94339475 – 16:94347709 Mb), located between *Vps26c* and *Dyrk1a* as previously described (*8*). The integrity of the Del4Tyb mutation was validated by comparative genome hybridization (Figure S4A). ES cells carrying the *Setd4*^tm1a(KOMP)Wtsi^ allele (*55*) were obtained from the International Knockout Mouse Consortium and used to establish a mouse strain which was bred first to Tg(CAG-Flpo)1Afst mice (*56*) which express Flp in the germline to delete LacZ and Neo genes, generating mice with the *Setd4*^tm1c(KOMP)Wtsi^ (*Setd4*^fl^) allele in which exon 6 (ENSMUSE00001268769) is flanked by loxP sites. These in turn were bred to Tg(Prm-cre)70Og mice (*57*) expressing Cre in the male germline to generate mice bearing the *Setd4*^tm1d(KOMP)Wtsi^ (*Setd4*^-^) allele in which exon 6 had been deleted. Mice with the *Mir802*^em1Tyb^ (*Mir802*^-^) allele were generated by direct injection of Cas9 and 2 guide RNAs (5’-TCTACATAACCTACCGACTGCGG-3’ and 5’-ACGCCCTCCGAGGACACCCCAGG-3’) into mouse zygotes to generate a 144 bp deletion (16:93166587 - 16:93166763) of the *Mir802* gene. Generation of mice with the *Dyrk1a*^tm2Yah^ (*Dyrk1a*^K188R^) allele will be described elsewhere (Y. Herault). Since both Dp1Tyb and *Dyrk1a*^+/-^ mice are poor breeders, we found it impossible to breed sufficient numbers of Dp1Tyb*Dyrk1a*^+/+/-^ mice (nomenclature indicates mice with two WT alleles of *Dyrk1a* and one deleted allele) by simply intercrossing these strains. To overcome this, rare mice bearing both the Dp1Tyb and *Dyrk1a*^-^ alleles were crossed to C57BL/6J mice and a pup was identified where a crossover had brought the two alleles onto the same chromosome. The resulting Dp1Tyb*Dyrk1a*^+/+/-^ mice bred well and were maintained as a separate strain, with WT littermate embryos used as controls. RNAseq analysis confirmed that one *Dyrk1a* allele had a deletion of exons 7 and 8 (Figure S4B). All mice were bred and maintained on a C57BL/6J background (backcrossed for ≥ 10 generations), initially at the MRC National Institute for Medical Research and then at the Francis Crick Institute. In all cases, littermate embryos were used as controls for mutant embryos. All animal experiments were carried out under the authority of a Project Licence granted by the UK Home Office, and were approved by the Animal Welfare Ethical Review Body of the Francis Crick Institute. Numbers of protein-coding genes in different mouse strains were determined using the Biomart function in Ensembl on mouse genome assembly GRCm39, filtering for protein-coding genes, excluding three genes: ENSMUSG00000116933 which is a partial transcript for *Atp5o* (ENSMUSG00000022956), *Gm49711*, which is an alternatively spliced form of *Mrps6*, and *Gm49948* which is a fusion transcript of some exons from *Igsf5* and *Pcp4*. Note that the numbers of duplicated coding genes in the Dp strains have changed since our original publication (*8*), due to changes in gene annotation.

### Array Comparative Genome Hybridization (aCGH)

Genomic DNA was prepared from tails of Del4Tyb (test) and C57BL/6J (control) mice. DNA, labeled with Cy3 (test) and Cy5 (control), was hybridized by Roche Diagnostics Limited to a mouse 3×720K array (Roche NimbleGen) containing 50-75mer probes designed based on mouse genome assembly MGSCv37. The hybridized aCGH slides were scanned in the Cy3 and Cy5 channels. The Log2 ratios of the test/control signals were calculated for all probes and the ratios for the probes on Mmu16 were plotted in genomic order to visualize the deleted region.

### RNA sequencing (RNAseq)

Frozen human fetal hearts (13-14 pcw, 5 DS and 5 euploid age- and sex-matched samples) were obtained from the MRC-Wellcome Trust Human Developmental Biology Resource (HDBR). Frozen tissues were placed in RLT lysis buffer (Qiagen) in gentleMACS M-tubes (Miltenyi Biotec), homogenized using the gentleMACS Octo dissociator (Miltenyi Biotec) and RNA was extracted using the RNAeasy Maxi kit (Qiagen). RNA integrity numbers (RIN) of the samples were 7-9.4. Stranded polyA-enriched libraries were made using the KAPA mRNA HyperPrep kit (Roche) according to the manufacturer’s instructions and sequenced on the HiSeq 4000 (Illumina) with single-ended 100 base reads. An average of 34 million reads were generated per sample.

E13.5 mouse hearts from Dp1Tyb, Dp1Tyb*Dyrk1a*^+/+/-^, Dp3Tyb and Ts1Rhr embryos and WT littermate embryos for each strain (5 per genotype) were dissected and snap frozen. Frozen tissues were placed in lysis buffer and homogenized using a cordless Pellet pestle (Sigma-Aldrich) followed by RNA extraction using RNAeasy mini kit (Qiagen). RNA integrity numbers (RIN) of the samples were 9.8-10. Stranded polyA-enriched libraries were made using the KAPA mRNA HyperPrep kit (Roche) according to the manufacturer’s instructions and sequenced on the HiSeq 2500 or HiSeq 4000 (Illumina) with single ended 100 base reads. An average of 24 million reads were generated for the Dp1Tyb and Dp1Tyb*Dyrk1a*^+/+/-^ and WT samples and an average of 34 million reads for Dp3Tyb, Ts1Rhr and corresponding WT samples. Adaptor trimming was performed with Trimmomatic/0.36-Java-1.7.0_80 with parameters “LEADING:3 TRAILING:3 SLIDINGWINDOW:4:20 MINLEN:36” (*58*). The RSEM package (v.1.2.31) in conjunction with the STAR alignment algorithm (v.2.5.2a) was used for the read mapping and gene-level quantification with respect to mouse genome assembly GRCm38 (release 86) for the mouse data sets and human genome assembly GRCh38 (release 89) for the human samples (*59, 60*). Unsupervised hierarchical clustering was carried out using Eisen distance with the heatmap.2 function from gplots (*61*). Differential expression analysis was performed with the DESeq2 package within the R programming environment (*62, 63*). The significance threshold for the identification of differentially expressed genes was set as an adjusted *P* value ≤0.05. Gene set enrichment analysis was carried out with the GSEA software (version 2.2.3) (*10*). The software compares ranked lists of genes, in this case, differentially expressed genes ranked by the “stat” value in decreasing order, with the following gene sets from the Molecular Signature Database: “c2.cp.v7.0.symbols.gmt”, “c5.bp.v7.0.symbols.gmt”, “h.all.v7.0.symbols.gmt” downloaded from the Broad Institute (http://www.broad.mit.edu/gsea/). GSEAPreranked was used with “classic” enrichment analysis not excluding large datasets.

For the pharmacological rescue experiment, 5 E13.5 hearts were analyzed for each of the 4 conditions: 2 genotypes (WT and Dp1Tyb) x 2 treatments (LCTB-21 and iso-LCTB-21). Both male and female embryos were analyzed, with the same numbers of each sex in each of the 4 groups. Samples were dissected and snap frozen. Frozen tissues were placed in lysis buffer and homogenized using a cordless Pellet pestle (Sigma-Aldrich) followed by RNA extraction using RNAeasy micro kit (Qiagen). RNA integrity numbers (RIN) of the samples were 10. Stranded polyA enriched libraries were generated using NEBNext^®^ Ultra^™^ II Directional RNA Library Prep Kit for Illumina according to manufacturer’s instructions. An average of 25 million paired end reads per library (PE100) were generated on a NovaSeq6000 (Illumina). All samples were processed using the nf-core RNAseq pipeline (v.3.10.1) operating on Nextflow (v.22.10.3) (*64, 65*). Settings for individual tools were left as standard for this workflow version unless otherwise specified. Alignment to GRCm38 (release 95) was performed using STAR (v2.7.10a) (*60*), followed by quantification via RSEM (v1.3.1) (*59*). Results were further processed in R (version 4.2.0) (*63*). Outputs of the RNAseq pipeline were assessed for quality using the metrics provided by the pipeline’s inbuilt quality control packages, PCA (package stats v4.2.0), and correlation analyses between samples (package stats v4.2.0). Differential expression analysis was performed with DESeq2 (v1.38.3) (*62*), using a model accounting for differences in litter, sex, treatment and genotype (*∼ litter + sex + treatment * genotype*). The significance threshold for the identification of differentially expressed genes was set as an adjusted *P* value ≤0.05, where *P* values were adjusted for false discovery rates according to the Independent Hypothesis Weighting method (package IHW, v1.26.0) (*66, 67*). Bulk RNAseq data have been deposited in the Gene Expression Omnibus, accession codes: GSE196447 and GSE239798.

### Single cell RNAseq

E13.5 mouse hearts from Dp1Tyb, Dp1Tyb*Dyrk1a*^+/+/-^ and WT littermate embryos were dissected and subsequently dissociated into single cells using the Neonatal Heart Dissociation kit and gentleMACS Octo dissociator (Miltenyi Biotec). Cells in suspension were methanol fixed, filtered through a 30 µm cell strainer, spun down and stored at −20°C. Fixed cells were resuspended in PBS, 0.04% BSA, assessed for viability using trypan blue staining and quantified using the EVE automated cell counter (Cambridge Bioscience). Cells were processed using the Chromium 3’ mRNA-Seq version 2 kit (10x Genomics). Samples were sequenced on the HiSeq 4000 acquiring at least 142,000 reads per cell, achieving a sequencing saturation of 73 - 90%.

Cell Ranger software (version 2.1.1, 10x Genomics) was used to de-multiplex Illumina BCL output, create fastq files and generate single cell feature counts for each library using GRCm38 (v1.2.0) as reference. All subsequent analyses were performed in R (v3.6.1) using the Seurat (v 3.1) package (*63, 68*). Genes were removed if they were expressed in 3 or fewer cells and cells with < 200 expressed genes detected were also removed. After quality control and filtering 3,384 cells were retained for further analysis. Data was integrated following Seurat’s vignette. In brief, for each sample the top 2000 most variable genes were selected for data integration using Canonical correlation Analysis (CCA) with 50 dimensions for dimensional reduction using tSNE and cluster calling using the Louvain algorithm. Clusters were visualized using the Uniform Manifold Approximation and Projection (UMAP). Upon initial examination of the clusters, we detected small clusters corresponding to red blood cells and lung tissue, both of which were removed. Subsequently data was normalized and integrated, and clustering analysis was repeated with the same parameters as above, with the addition that the effects of cell cycle were regressed out. A cluster resolution of 0.4 was used to define clusters for further analysis. Cluster markers were identified using “FindConservedMarkers” with default parameters. Differentially expressed genes within each cluster were identified using the DESeq2 test in “FindMarkers” using data in which the effects of cell cycle had not been regressed out. scRNAseq data have been deposited in the Gene Expression Omnibus, accession code: GSE196447.

### Proteomics

E13.5 mouse hearts from Dp1Tyb*Dyrk1a*^+/+/-^ and WT littermate embryos (5 per genotype) were dissected, snap frozen and stored at −80°C. Frozen tissues were placed in 8M Urea with phosSTOP protease inhibitors (Roche) and homogenized using a cordless Pellet pestle (Sigma-Aldrich). Lysates were reduced, alkylated and digested with tryspin using standard methods. Nest Group C_18_ MacroSpin columns (SMMSS18V) were used to concentrate and clean up the peptides according to the manufacturer supplied instructions, prior to labeling with TMT10plex Isobaric Label Reagent Set (ThermoFisher Scientific). TMT-labeled samples were pooled, enriched for phospho-peptides using the High-Select Fe-NTA phosphopeptide enrichment kit (A32992, ThermoFisher Scientific) and using Titansphere TiO 5 μm bulk media (GL Sciences) according to the manufacturer supplied protocols. Samples were analyzed by LC-MS/MS on an Orbitrap Fusion Lumos Mass Spectrometer (ThermoFisher Scientific). Raw data were processed in MaxQuant v1.6.0.13 (https://www.maxquant.org/), with the database search conducted against the canonical sequences of the UniProt Mus Musculus complete proteome, downloaded August 2017. Statistical analysis was carried out using Perseus (https://www.maxquant.org/perseus/). In brief, abundance values for each protein were log2 transformed, median normalized and a Welch t-test was used to evaluate the significance of differences between samples of the two genotypes, generating an FDR corrected p-value and log2(fold change of the geometric means). The mass spectrometry proteomics data have been deposited with the ProteomeXchange Consortium (http://proteomecentral.proteomexchange.org) via the PRIDE partner repository (*69*), dataset identifier PXD013053.

### HREM imaging and 3D modeling

E14.5 embryonic hearts were dissected and fixed for 30 min in 4% paraformaldehyde followed by a 1 h wash in distilled water and fixed a second time overnight. Fixed samples were dehydrated and embedded in modified JB4 methacrylate resin (*70*). Samples were sectioned at 2 μm and imaged using a Jenoptik camera with an isometric resolution of 2 μm. Data sets were normalized and subsampled prior to 3D volume rendering using OsiriX MD (*71*). Phenotype analysis was performed blind for genotype and classification of type of CHD was carried out as previously described (*72*). Mutant and control WT hearts were taken from the same litters.

### Flow cytometry

E13.5 mouse hearts from Dp1Tyb and WT littermate embryos and Dp1Tyb*Dyrk1a*^+/+/-^ and their WT littermates were dissociated into single cells using the Neonatal Heart Dissociation kit and gentleMACS Octo dissociator (Miltenyi Biotec). To evaluate mitochondrial membrane potential and content cells in suspension were incubated with 20 nM tetramethylrhodamine methyl ester (TMRM, Thermo Fisher) and 20 nM of Mitotracker Green (MTG, Thermo Fisher), respectively, for 30 min at 37°C. Cells were then stained with anti-CD106-APC, anti-CD31-BV785 and Zombie Aqua (all Biolegend). CD106 (VCAM-1) is predominantly expressed on cardiomyocytes in mouse embryonic hearts (*73*). CD31 (PECAM1) is expressed on both endocardial and endothelial cells, but in E13.5 hearts most CD31^+^ cells are endocardial cells (Figure 2C). Zombie Aqua was used to distinguish live and dead cells. To analyze cell cycle phases, dissected E13.5 mouse hearts were incubated with 10µM EdU at 37°C for 30 min, and then dissociated into single cells using the Neonatal Heart Dissociation kit and gentleMACS Octo dissociator (Miltenyi Biotec). Cells in suspension were stained in PBS with anti-CD106-APC, anti-CD31-BV785 (Biolegend) and Live/dead Near-IR (ThermoFisher) for 30 min, followed by fixation in 4% Paraformaldehyde (PFA) for 20 min. EdU was detected using the Click-it EdU kit (Life Technologies), and DNA was stained with FxCycle violet (ThermoFisher). Data were acquired on an LSR Fortessa X-20 cell analyzer (BD) and analyzed using FlowJo v9.

### Immunoblot analysis

E13.5 mouse hearts from Dp1Tyb and WT littermate embryos were dissected and snap frozen. Frozen tissues were placed in RIPA lysis buffer and homogenized using a cordless Pellet pestle (Sigma-Aldrich) followed by protein extraction. Total protein lysates (20 μg) were resolved on a denaturing 4-10% precast SDS-PAGE (Bio-Rad) and probed with rabbit monoclonal antibodies against phospho-RB1 (Ser807/Ser811) and GAPDH (Cell Signaling Technology, 8516 and 5174, respectively). Binding of primary antibodies was detected using AF680–conjugated anti-rabbit IgG (Thermo Fisher Scientific). Fluorescence from the secondary reagents was detected using an Odyssey (LI-COR Biosciences). For quantitation, signal from phospho-RB1 was normalized to GAPDH.

### Metabolic analysis

E13.5 mouse hearts from Dp1Tyb and WT littermate embryos and Dp1Tyb*Dyrk1a*^+/+/-^ and their WT littermates were dissociated into single cells as described above. To measure rates of mitochondrial respiration, cells were seeded onto Seahorse 8-well plates in in XF D-MEM Medium pH 7.4 containing 10 mM glucose, 2 mM L-glutamine and 1 mM sodium pyruvate (all Agilent Technologies). The plates were centrifuged at 100xg for 3 min, kept at 37°C for 45-60 min and the oxygen consumption rate (OCR) was measured using a Seahorse XFp Analyzer (Agilent Technologies). To investigate mitochondrial respiration phenotypes, a Seahorse XFp Cell Mito Stress Test kit (Agilent Technologies) was used. Cells were analyzed for 20 min and then oligomycin (1 µM, final concentration) was added, followed by carbonyl cyanide 4-(trifluoromethoxy) phenylhydrazone (FCCP) (1 µM, final concentration) 20 min later, and rotenone and antimycin A (0.5 µM, final concentration) a further 20 min after that. To measure glycolysis, cells were seeded onto Seahorse 96-well plates in in XF D-MEM Medium pH 7.4 containing 2 mM L-glutamine. The plates were centrifuged at 100xg for 3 min, kept at 37°C for 45-60 min and the extra-cellular acidification rate (ECAR) was measured using a Seahorse XFe96 Analyzer, using a Seahorse XFe96 Glycolysis stress Test kit (Agilent Technologies). To establish the basal ECAR level, ECAR was measured for 20 min at the beginning of the assay. After 20 min, glucose (10 mM) was added, followed by oligomycin (1 μM) 20 min later, and 2-deoxy-D-glucose (50 mM) a further 20 min after that. Cells were fixed with 4% PFA for 15 min, nuclei were stained with 4′,6-diamidino-2-phenylindole (DAPI) and imaged using an EVOS microscope (Thermo Fisher). The Analyze Particles command from ImageJ was used to calculate numbers of nuclei which were used to normalize the OCR and ECAR data to numbers of cells in a well. Further analysis of OCR and ECAR was carried out using WAVE (version 2.6.1, Agilent Technologies).

### Mitochondrial shape analysis

E13.5 mouse hearts from Dp1Tyb and WT littermate embryos and Dp1Tyb*Dyrk1a*^+/+/-^ and their WT littermates were dissociated as previously described. Cells were seeded on 96-well plate (Greiner Bio-One) and cultured overnight to allow them to adhere. Mitochondria were stained by incubating the cells with 20nM of MitoTracker Deep Red (MTDR) for 30 min at 37°C and fixed with 2% PFA for 20 min at 37°C. Cells were stained with rat anti-mouse CD106 (BD Bioscience) overnight at 4°C, followed by goat anti-rat IgG Alexa Fluor 488 (Thermo Fisher) for 1 h at room temperature, and DAPI for 10 min at room temperature. Cells were imaged with the Opera Phenix High-Content Screening System (PerkinElmer). Initially, images were acquired using PreciScan imaging with a x20/NA 0.4 air lens and the locations of single cardiomyocytes (CD106^+^) were defined using Harmony software V4.9. Next, single cardiomyocytes were re-imaged using a ×63/NA 1.15 water-immersion lens. Z-stacks from 0 to 3 µm with a step size of 1 µm were acquired using excitation lasers at 405 nm (DAPI), 488 nm (CD106) and 640 nm (MTDR). The mitochondrial network was identified using the MTDR signal (mitochondria) transformed into Ridge texture using the SER (Saddles, Edges, Ridges) feature of Harmony V4.9 (PerkinElmer). Mitochondrial shape was assessed using aspect ratio (major axis length/minor axis length), a measure of the length to width ratio, and form factor (perimeter^2^/4ρχ[area]), reflecting the complexity and branching of mitochondria (*74*). These parameters were determined using Harmony software on maximum projection images (Figure S3A, B).

### Hypoxia analysis

E13.5 pregnant mice were injected i.p. with 60 mg/kg of Hypoxyprobe^TM^-1 (pimonidazole HCl, Hypoxyprobe, Inc.) in saline. Embryonic hearts were dissected 2 to 3 h post injection and fixed for 15 min in 4% paraformaldehyde followed by 1 h wash in distilled water and fixed a second time overnight. Hearts were paraffin-embedded and sectioned at 4 µm thickness and mounted on SuperFrost Plus slides. Hypoxyprobe was detected using a rabbit anti-pimonidazole antibody from the Hypoxyprobe Omni-Kit (Hypoxyprobe, Inc.), mouse anti-cardiac troponin (MA5-12960, Thermo Scientific) and rat anti-endomucin (Sc-65495, Santa Cruz) overnight at 4°C, followed by goat anti-rabbit IgG Cy3, goat anti-mouse IgG AF647 and goat anti-rat IgG AF488 (Thermo Fisher) for 1 h at room temperature, and DAPI for 10 min at room temperature.

### Treatment of pregnant mice with Leucettinib-21

C57BL/6J female mice were mated with C57BL/6J males. From 5 days post coitum (E5.5), pregnant mice were orally gavaged daily with 0.3, 3 or 30 mg/kg of Leucettinib-21 (Figure S5A, Perha Pharmaceuticals) in 0.5% carboxymethylcellulose sodium salt (Sigma 9004-32-4) until E14.5. Embryos were dissected 2 h after the final gavage, snap frozen and then analyzed by mass spectrometry for the presence of Leucettinib-21 (Oncodesign Services). Alternatively, C57BL/6J female mice were mated with Dp1Tyb males and pregnant mice were treated with 30 mg/kg of Leucettinib-21 or iso-Leucettinib-21 in 0.5% carboxymethylcellulose daily by oral gavage from E5.5 to E13.5 (for RNAseq) or E14.5 (for HREM). Embryonic hearts were dissected 2h after the final gavage and processed for RNAseq and HREM imaging as described above.

## Statistical analysis

Statistical analysis of RNAseq and proteomic data is described above. Other data were analyzed using Fisher’s exact test, Mann Whitney test or Kruskal-Wallis test.

## Acknowledgments

We thank James Turner for critical reading of this manuscript. We thank Fabrice Prin for help with HREM and Michael Howell for help with high throughput analysis of mitochondrial morphology. We thank the Advanced Sequencing, Proteomics, High Throughput Screening, Flow Cytometry, Advanced Light Microscopy, Experimental Histopathology, Genetic Manipulation and Biological Research Facilities of the Francis Crick Institute for sequencing, proteomics, mitochondrial morphology analysis, flow cytometry, imaging, histology, generation of genetically altered mice and for animal husbandry. We thank Mariona Arbones and Christian Lüscher for mouse strains.

## Author Contributions

Conceptualization: EMCF, VLJT

Data curation: EL-E, RA, ML

Formal analysis: EL-E, RA, ML, HF

Funding acquisition: EMCF, VLJT

Investigation: EL-E, RA, ML, DG, CB, HF, SW-S, MHH, DH, O-RS

Methodology: EL-E, RA, SW-S

Project administration: VLJT

Resources: EL-E, SW-S, VB, YH, ED, LM

Supervision: APS, MG, EMCF, VLJT

Visualization: EL-E, RA, ML

Writing – original draft: EL-E, RA, VLJT

Writing – review & editing: EL-E, RA, EMCF, VLJT

## Data and materials availability

All bulk and single-cell RNAseq data have been deposited in the Gene Expression Omnibus, accession code: GSE196447. Mass spectrometry proteomic data have been deposited in the ProteomeXchange via PRIDE, dataset identifier: PXD013053.

## Supplementary Material

### Supplementary Figures

**Figure S1.**
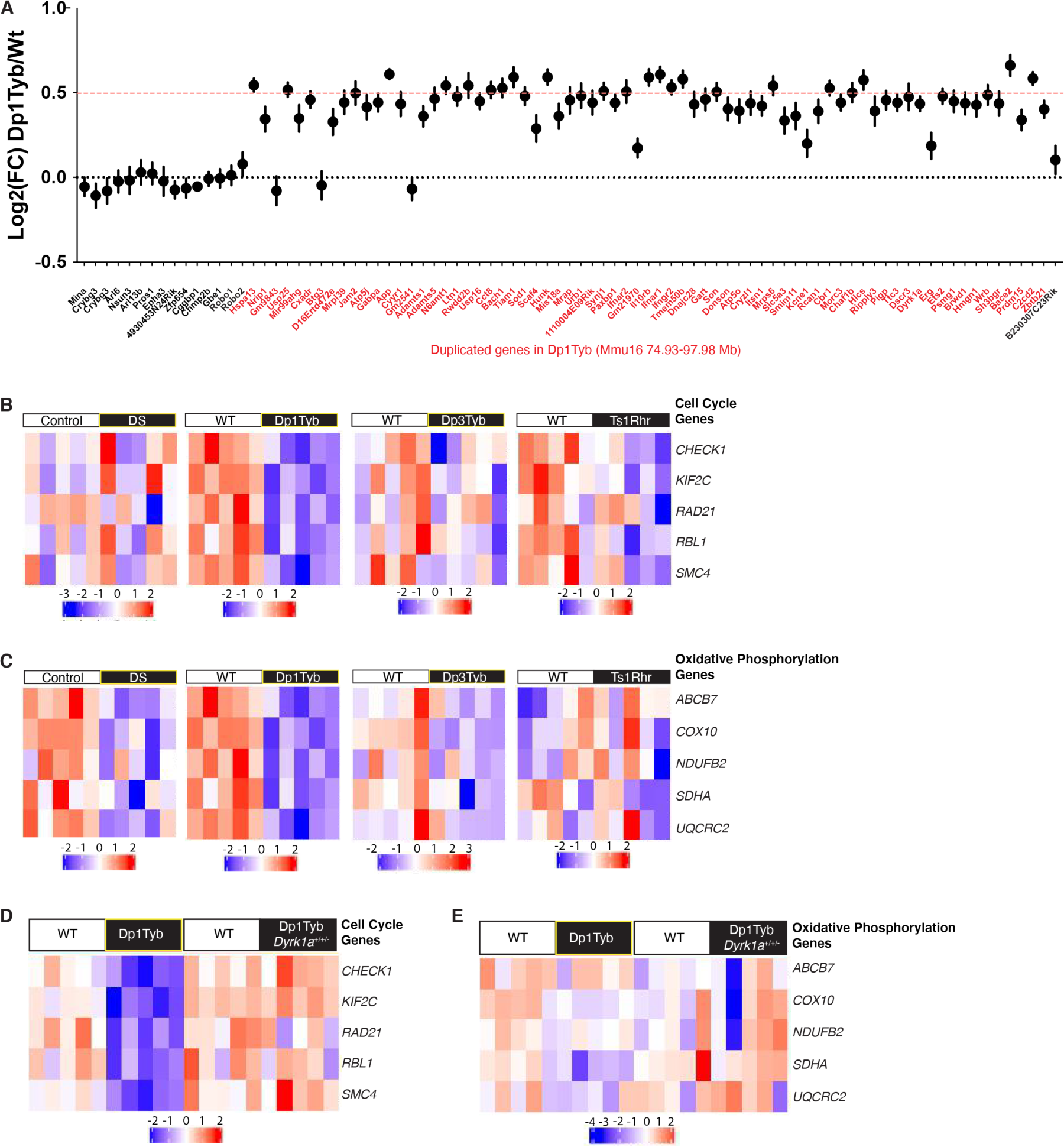
Altered gene expression in DS and Dp1Tyb embryonic hearts. (**A**) Mean±SEM log_2_(fold-change) of gene expression between WT and Dp1Tyb E13.5 hearts. Only expressed genes are shown, defined as genes whose mean expression is >5 TPM and having a measured p-value in DEseq2. Genes within the duplicated region of Mmu16 are shown in red. For comparison, 15 genes that are centromeric to this region of Mmu16 and one gene telomeric to this region and not duplicated are shown in black. Dashed red line indicates a fold change of 1.5 expected by the increased dosage of the duplicated genes; dotted black line represents no change in expression. All but three of the duplicated genes show increased expression. FC, fold-change. (**B-E**) Heatmaps showing change in gene expression of selected genes from (B, D) the Reactome cell cycle and (C, E) Hallmark oxidative phosphorylation gene sets in (B, C) DS and control euploid human embryonic hearts, and in Dp1Tyb, Dp3Tyb and Ts1Rhr mouse E13.5 hearts and their corresponding WT controls and (D, E) in Dp1Tyb and Dp1Tyb*Dyrk1a*^+/+/-^ embryonic hearts and their corresponding WT littermate controls. Red and blue colors indicate increased or decreased expression of indicated genes relative to the mean expression of each row using normalized log_2_ expression values.

**Figure S2.**
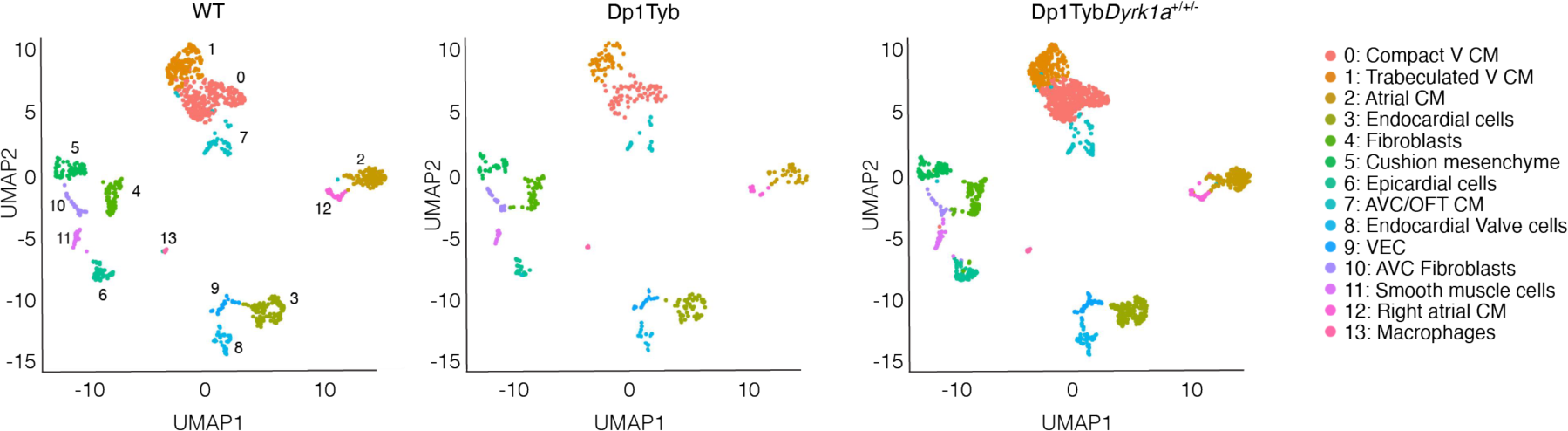
scRNAseq analysis of Dp1Tyb and Dp1Tyb *Dyrk1a*^+/+/-^ embryonic hearts. UMAP plots of scRNAseq data from WT, Dp1Tyb and Dp1Tyb *Dyrk1a*^+/+/-^ mouse E13.5 embryonic hearts. Clusters were generated by pooling data from all hearts and all genotypes as shown in Figure 2A. The plots in this figure use the same clustering but show cells in each genotype separately. Sample numbers: n=2 WT, 1 Dp1Tyb, 2 Dp1Tyb*Dyrk1a*^+/+/-^.

**Figure S3.**
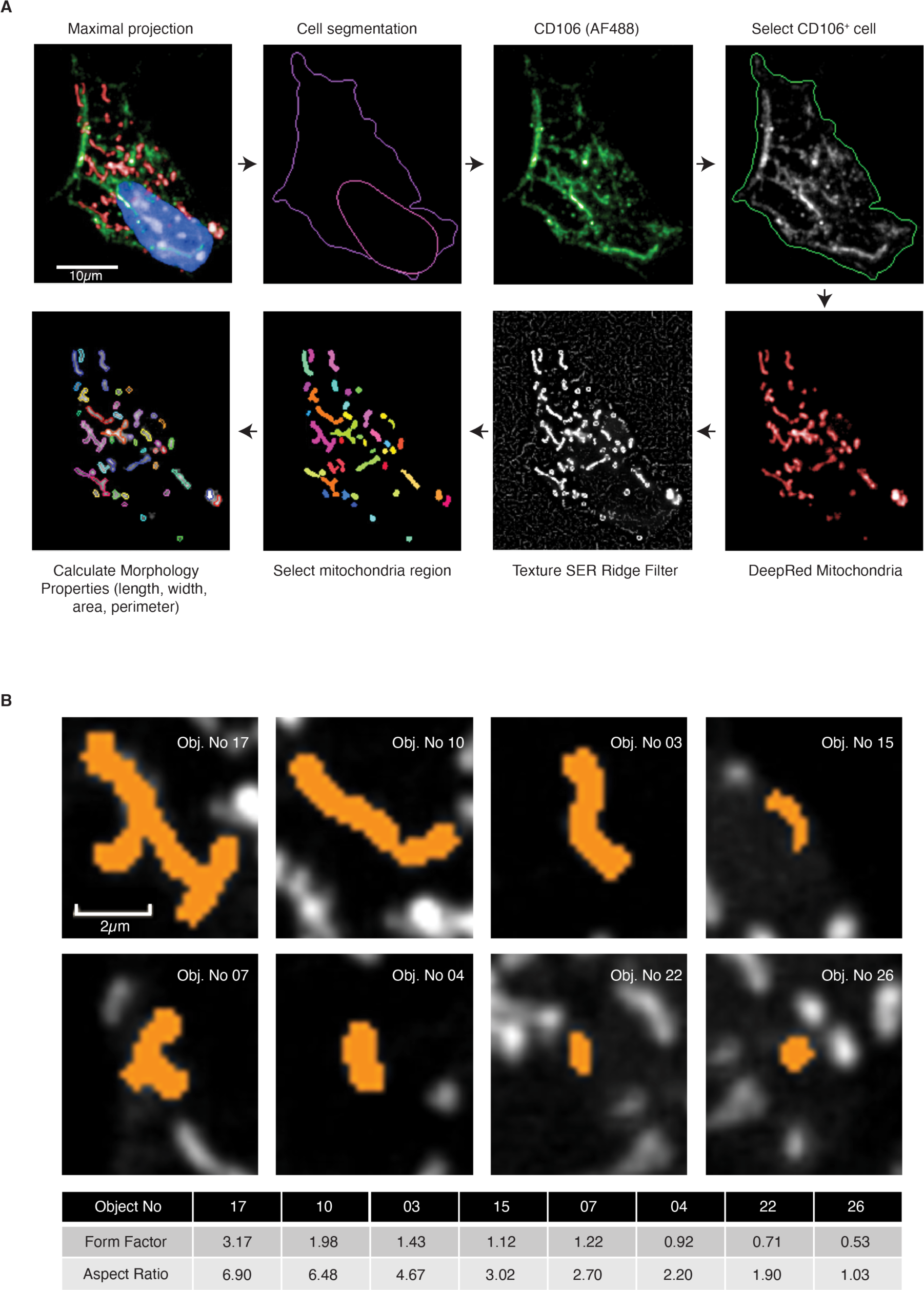
Quantitation of mitochondrial morphology. (**A**) Mouse embryonic heart single cells stained with MitoTracker Deep Red (MTDR), anti-CD106 (AF488) and DAPI were imaged by confocal microscopy. Maximal projection images (Z-stacks from 0 to 3 µm with a step size of 1 µm) were segmented to identify location of cells and nuclei. CD106^+^ (cardiomyocytes) cells were selected for further analysis. The mitochondrial network was identified using the MTDR signal (mitochondria) transformed into Ridge texture using the SER (Saddles, Edges, Ridges) feature of Harmony. Mitochondrial network morphology was assessed by measuring the aspect ratio (major axis length/minor axis length) and form factor (perimeter^2^/4ρχ[area]), which are measures of distortion from circularity and degree of branching, respectively. (**B**) Eight example images of mitochondria (objects, Obj.) showing their aspect ratios and form factors.

**Figure S4.**
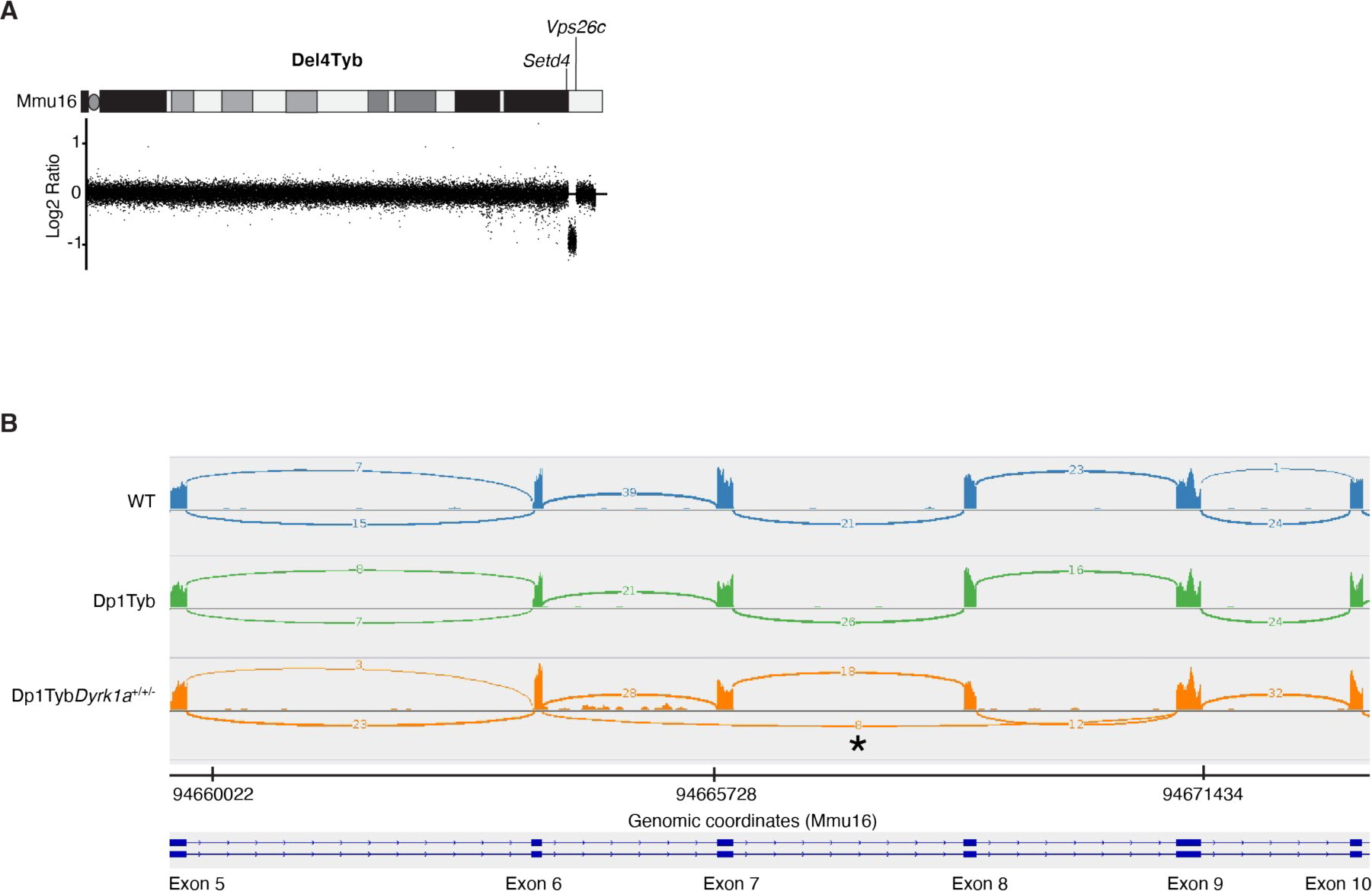
Validation of the Del4Tyb and Dp1Tyb*Dyrk1a*^+/+/-^ mouse strains. (**A**) Comparative genome hybridization analysis of the Del4Tyb mouse strain. Graph shows the log_2_(ratio) of the hybridization signal between Del4Tyb and C57BL/6J control mice. Each dot is a different probe along the length of Mmu16, in genomic order. Diagram at top shows a map of Mmu16 indicating the acrocentric centromere and the location of the *Setd4* and *Vps26c* genes marking the ends of the deletion. The deleted region is expected to have 0.5-fold decrease in DNA content, resulting in a log_2_(ratio) = −1. (**B**) Plot showing RNAseq reads mapped against the mouse genome within the *Dyrk1a* gene (exons 5 to 10) for E13.5 hearts from a WT, Dp1Tyb and Dp1Tyb*Dyrk1a*^+/+/-^ embryo. Reads can be seen mapping against each exon and arcs indicate reads spanning introns. Numbers on the arcs show the numbers of reads showing the given splicing event. The Dp1Tyb*Dyrk1a*^+/+/-^ embryo shows some reads connecting exon 6 to exon 9 (asterisk), as would be predicted if exons 7 and 8 had been deleted in one *Dyrk1a* allele. These connecting reads are not seen in WT or Dp1Tyb mice, confirming successful deletion of exons 7 and 8 in Dp1Tyb*Dyrk1a*^+/+/-^ embryos.

**Figure S5.**
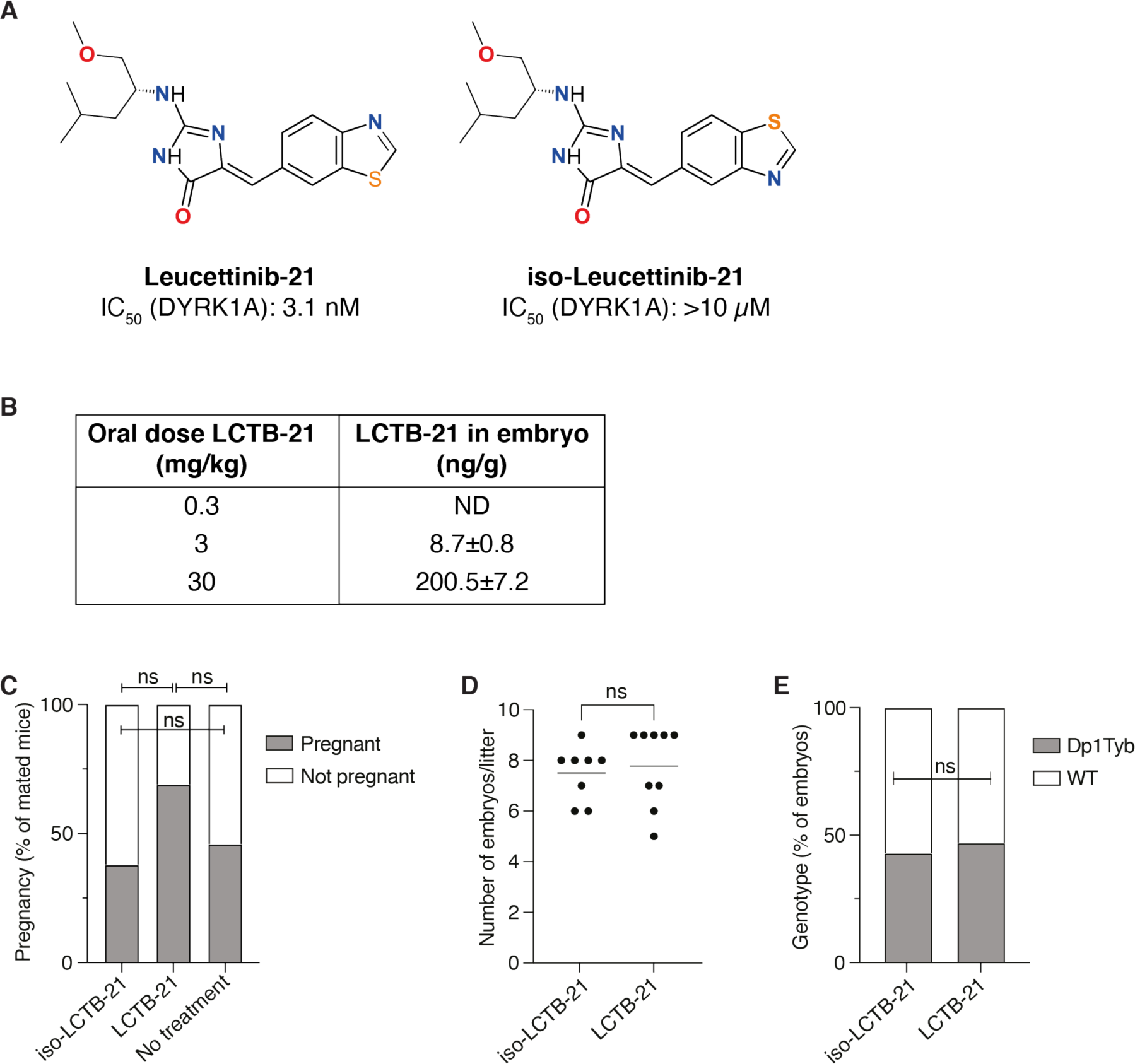
Treatment of pregnant mice with Leucettinib-21. (**A**) Chemical structure of Leucettinib-21 (LCTB-21) and iso-Leucettinib-21 (iso-LCTB-21) indicating IC_50_ concentrations for the inhibition of DYRK1A in vitro (LCTB-21 is compound 106 in *24*). (**B**) Pregnant C57BL/6J mice were treated with the indicated dose of LCTB-21 daily by oral gavage from E5.5 to E14.5 and embryos analyzed for levels of LCTB-21 by mass spectrometry, with values listed as mean (±SEM) ng of LCTB-21 per g of embryo tissue. ND, not detected. (**C-E**) C57BL/6J female mice that had successfully mated with Dp1Tyb male mice as judged by the presence of a vaginal plug on the day following mating (embryonic day 0.5, E0.5) were treated daily with LCTB-21 or iso-LCTB-21 by oral gavage from E5.5 to E14.5 (Figure 8A). Graphs show percentage of mated mice that (C) were pregnant at E14.5, (D) number of embryos per litter and (E) percentage of embryos whose genotype was WT or Dp1Tyb. Statistical analysis using Fishers exact (C, E) and Mann Whitney (D) tests; ns, not significant. Sample numbers: B, n = 8 (0.3 or 3 mg/kg) and 4 (30 mg/kg) embryos; C, n = 30 (iso-LCTB-21), 19 (LCTB-21) and 24 (no treatment) mice; D, n = 8 (iso-LCTB-21) and 9 (LCTB-21) litters; E, n = 60 (iso-LCTB-21) and 70 (LCTB-21) embryos.

**Figure S6.**
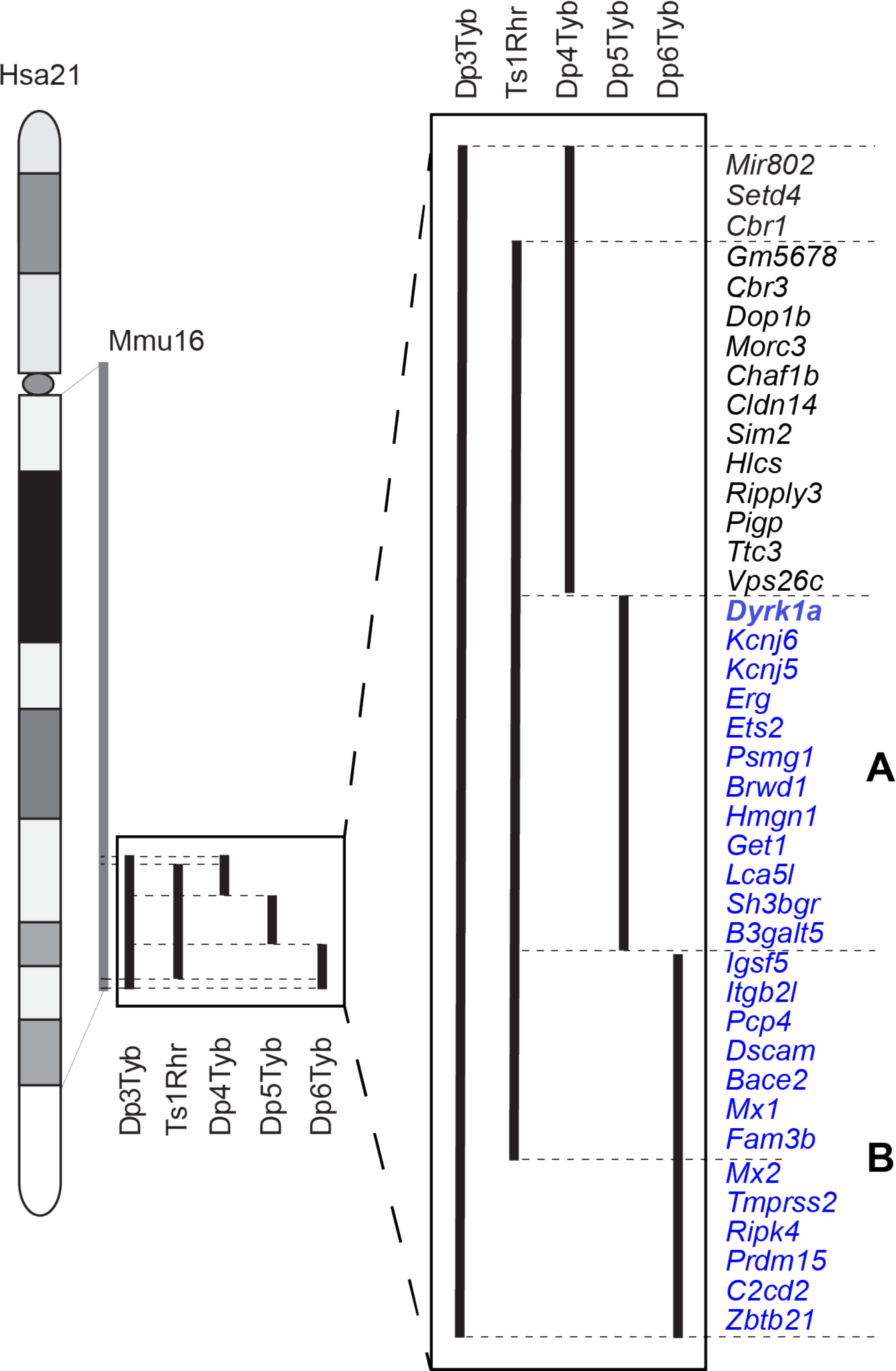
Regions of Hsa21 containing dosage-sensitive genes causing CHD. Diagram of Hsa21 showing main cytogenic regions (rectangles of different shades) and the centromere (oval). Grey bar indicates the Hsa21-orthologous region of Mmu16. Black lines indicate regions of Mmu16 duplicated in the Dp3Tyb, Ts1Rhr, Dp4Tyb, Dp5Tyb and Dp6Tyb mouse strains. These regions are expanded showing all known protein coding genes within them and one microRNA gene (*Mir802*). Dp3Tyb mice have CHD, but Ts1Rhr, Dp4Tyb, Dp5Tyb and Dp6Tyb do not, implying that there must be at least two causative genes. The genes in the Dp4Tyb region are not required for CHD, since a cross of Del4Tyb to Dp1Tyb did not affect the frequency of CHD. This implies that the two causative genes must lie in the regions duplicated in Dp5Tyb and Dp6Tyb respectively (regions A and B, genes in blue). We term this the 2-locus hypothesis. The *Dyrk1a* gene (bold) is required in three copies for the CHD phenotype, but there may be other genes in region A that are also required. The second unknown gene lies in region B and is most likely one of the 6 distal genes present in 3 copies in Dp3Tyb but not Ts1Rhr mice (*Mx2*-*Zbtb21*). Genes in regions that do not cause CHD are in black.

### Supplementary Tables

**Table S1. Human fetal hearts used for RNAseq analysis.**

List of 10 human fetal heart samples obtained from HDBR (5 DS, 5 euploid) which were used for RNAseq showing gestational age and sex.

**Table S2. RNAseq analysis of human Down Syndrome fetal hearts.**

Sheet 1: Read me. Sheet 2: Expression of all genes in all samples in transcripts per million reads (TPM), and mean expression in each of the two genotypes (DS and euploid). Sheet 3: Differential gene expression analysis from DESeq2 showing for each gene its location, mean expression across all samples, log2(fold change) between DS and euploid and the *p*-value and adjusted *p*-value for this difference.

**Table S3. RNAseq analysis of Dp1Tyb E13.5 embryonic hearts.**

Sheet 1: Read me. Sheet 2: Expression of all genes in all samples in transcripts per million reads (TPM), and mean expression in each of the two genotypes (Dp1Tyb and WT). Sheet 3: Differential gene expression analysis from DESeq2 showing for each gene its location, mean expression across all samples, log2(fold change) between Dp1Tyb and WT and the *p*-value and adjusted *p*-value for this difference.

**Table S4. RNAseq analysis of Dp3Tyb E13.5 embryonic hearts.**

Sheet 1: Read me. Sheet 2: Expression of all genes in all samples in transcripts per million reads (TPM), and mean expression in each of the two genotypes (Dp3Tyb and WT). Sheet 3: Differential gene expression analysis from DESeq2 showing for each gene its location, mean expression across all samples, log2(fold change) between Dp3Tyb and WT and the *p*-value and adjusted *p*-value for this difference.

**Table S5. RNAseq analysis of Ts1Rhr E13.5 embryonic hearts.**

Sheet 1: Read me. Sheet 2: Expression of all genes in all samples in transcripts per million reads (TPM), and mean expression in each of the two genotypes (Ts1Rhr and WT). Sheet 3: Differential gene expression analysis from DESeq2 showing for each gene its location, mean expression across all samples, log2(fold change) between Ts1Rhr and WT and the *p*-value and adjusted *p*-value for this difference.

**Table S6. Expression of E2F targets and Hypoxia gene sets**

Gene set enrichment analysis (GSEA) of the Hallmark gene set of E2F targets and Hypoxia gene sets. Sheet 1: Read me. Sheet 2: GSEA of E2F target genes in the comparison of transcriptomes of DS and euploid fetal hearts, identifying genes in the leading edge. Sheet 3: GSEA of E2F target genes in the comparison of transcriptomes of Dp1Tyb and WT embryonic hearts, identifying genes in the leading edge. Sheet 4: GSEA of E2F target genes in the comparison of transcriptomes of Dp1Tyb*Dyrk1a*^+/+/-^ and WT embryonic hearts, identifying genes in the leading edge. Sheet 5: Comparison of the leading genes in the GSEA of E2F target genes of human DS v euploid and mouse Dp1Tyb v WT embryonic hearts. Sheet 6: GSEA of Hypoxia genes in the comparison of transcriptomes of Dp1Tyb and WT embryonic hearts.

**Table S7. RNAseq analysis of Dp1Tyb*Dyrk1a*^+/+/-^ E13.5 embryonic hearts.**

Sheet 1: Read me. Sheet 2: Expression of all genes in all samples in transcripts per million reads (TPM), and mean expression in each of the two genotypes (Dp1Tyb*Dyrk1a*^+/+/-^ and WT). Sheet 3: Differential gene expression analysis from DESeq2 showing for each gene its location, mean expression across all samples, log2(fold change) between Dp1Tyb*Dyrk1a*^+/+/-^ and WT and the *p*-value and adjusted *p*-value for this difference.

**Table S8. Proteomic analysis of Dp1Tyb*Dyrk1a*^+/+/-^ E13.5 embryonic hearts.**

Sheet 1: Read me. Sheet 2: Abundance of phospho-peptides in all samples, the Welch difference, a measure of fold-change, between Dp1Tyb*Dyrk1a*^+/+/-^ and Dp1Tyb hearts, and the *p*-value. Sheet 3: Abundance of peptides in all samples, the Welch difference, a measure of fold-change, between Dp1Tyb*Dyrk1a*^+/+/-^ and Dp1Tyb hearts, and the *p*-value.

**Table S9. Effect of Dyrk1a dosage on differential gene expression.**

Sheet 1: Read me. Sheet 2: DESeq2 analysis of the transcriptomes of Dp1Tyb v WT embryonic hearts. Sheet 3: DESeq2 analysis of the transcriptomes of Dp1Tyb v Dp1Tyb*Dyrk1a*^+/+/-^ embryonic hearts. Sheet 4: Differentially expressed genes in common between those identified in sheets 2 and 3. Sheet 5: STRING analysis of the common upregulated genes. Sheet 6: STRING analysis of the common downregulated genes.

**Table S10. RNAseq of Dp1Tyb and WT embryonic hearts from pregnant mice treated with Leucettinib-21 or iso-Leucettinib-21.**

Sheet 1: Read me. Sheet 2: Normalized counts of all genes in all samples, and mean expression for each condition. Sheet 3: Differential gene expression analysis from DESeq2 showing for each gene its location, mean expression across all samples, log2(fold change) between Dp1Tyb embryonic hearts treated with iso-Leucettinib-21 and WT embryonic hearts treated with iso-Leucettinib-21, the *p*-value and an indication if the adjusted p-value reached significance. Sheet 4: Differential gene expression analysis from DESeq2 showing for each gene its location, mean expression across all samples, log2(fold change) between Dp1Tyb embryonic hearts treated with Leucettinib-21 and WT embryonic hearts treated with iso-Leucettinib-21, the *p*-value and an indication if the adjusted p-value reached significance.

